# NAD^+^ repletion rescues female fertility during reproductive ageing

**DOI:** 10.1101/721985

**Authors:** Michael J. Bertoldo, Dave R. Listijono, Wing-Hong Jonathan Ho, Angelique H. Riepsamen, Xing L. Jin, Kaisa Selesniemi, Dale M. Goss, Saabah Mahbub, Jared M. Campbell, Abbas Habibalahi, Wei-Guo Nicholas Loh, Neil A. Youngson, Jayanthi Maniam, Ashley S.A. Wong, Dulama Richani, Catherine Li, Yiqing Zhao, Maria Marinova, Lynn-Jee Kim, Laurin Lau, Rachael M Wu, A. Stefanie Mikolaizak, Toshiyuki Araki, David G. Le Couteur, Nigel Turner, Margaret J. Morris, Kirsty A. Walters, Ewa Goldys, Christopher O’Neill, Robert B. Gilchrist, David A. Sinclair, Hayden A. Homer, Lindsay E. Wu

**Author notes:** These authors contributed equally to this work (Michael J. Bertoldo, Dave R. Listijono, Wing-Hong Jonathan Ho, David A. Sinclair, Hayden A. Homer, Lindsay E. Wu).

## Abstract

Female infertility is a common and devastating condition with life-long health, emotional and social consequences. There is currently no pharmacological therapy for preserving oocyte quality during aging, which is the strongest risk factor for infertility. This leads to an age dependent decline in natural conception and IVF success rates (1). Here, we show that this is due in part to declining levels of the metabolic cofactor nicotinamide adenine dinucleotide (NAD^+^), and that restoring NAD^+^ levels with its metabolic precursor nicotinamide mononucleotide (NMN) rejuvenates oocyte quality and quantity in aged animals, leading to improved fertility. These benefits extend to the developing embryo, where NMN supplementation in embryo culture media following IVF enhances blastocyst formation in older mice. The NAD^+^ dependent deacylase SIRT2 is sufficient, but not essential, to recapitulate the benefits of *in vivo* NMN treatment, and transgenic overexpression of SIRT2 maintains oocyte spindle assembly, accurate chromosome segregation, decreased oxidative stress and overall fertility with ageing. Pharmacological elevation of NAD^+^ may be an effective, non-invasive strategy for restoring and maintaining female fertility during ageing, and for improving the success of IVF.

## Introduction

Across the developed world, there is a trend for delaying pregnancy until later in life for socio-economic reasons (2). This older age of parenthood is increasingly conflicting with age-dependent barriers to female fertility (1, 3), leading to increased demand for infertility treatments, including assisted reproduction technologies (ART) such as *in vitro* fertilisation (IVF). Resorting to this procedure has several disadvantages, as it is invasive, carries health risks (4), is expensive and has a limited success rate. Repeated IVF failures are a substantial source of emotional distress, and failure to conceive offspring is a substantial source of relationship breakdown (5).

The rate-limiting factors for successful pregnancies in IVF are oocyte quantity and quality, both of which start to decline from the middle of the third decade of life in humans (1, 3). Despite the enormous need, there are no clinically viable strategies to either preserve or rejuvenate oocyte quantity or quality during ageing. There is a major need in reproductive medicine for a non-invasive, pharmacological treatment to maintain or restore oocyte quantity and/or quality during ageing. The effect of such a therapy would be to alleviate a rate-limiting barrier to IVF success, or increase the chances of unaided conception, without having to resort to IVF.

The molecular basis for the decline in oocyte quality with advancing age is not clear but is certainly multifactorial. The key factors thought to be involved include genome instability, reduced mitochondrial bioenergetics, increased reactive oxygen species (ROS), and impaired fidelity during meiotic chromosome segregation due to disrupted spindle assembly and compromised function of the spindle assembly checkpoint (SAC) surveillance system (6). This latter hypothesis is evidenced by an increased rate of aneuploidy in embryos with increased maternal age (7), and the increased incidence of offspring born with chromosomal abnormalities such as Trisomy 21 (8), which causes Down’s Syndrome. The molecular cause of chromosome mis-segregation in oocytes with advancing age is still unknown, and as a result there are no pharmacological strategies to correct this problem. Understanding the molecular basis of this defect could lead to therapies which could maintain, or even rescue female fertility with advancing age.

Molecular defects in oocytes during ageing closely resemble the pathophysiology of somatic cell ageing, which is similarly characterised by epigenetic dysfunction and genome instability (9), declining mitochondrial bioenergetics (10), and impaired chromosome segregation leading to senescence (11). Given these common features, reproductive decline could be viewed through the same lens as somatic ageing. While somatic tissues undergo continual renewal through turnover of cells that can be replaced by a self-renewing population of resident precursor stem cells, primordial follicles in the ovary are laid down during *in utero* development in humans, where they form a finite pool that does not undergo self-renewal and cannot be replenished. Oocytes are therefore the oldest cells in the body: given this, it is unsurprising that while the somatic cells of a 35 year old female will have turned over many times in her lifetime, some being only days old, oocytes that were formed 35 years earlier in primordial follicles during fetal development will be highly susceptible to age-related dysfunction.

One key mechanism of ageing that has come to light in recent years is the role of nicotinamide adenine dinuclueotide (NAD^+^), a prominent redox cofactor and enzyme substrate that is essential to driving the electron transport chain for oxidative phosphorylation, and for other catabolic processes throughout the cell. NAD^+^ is also used as a substrate for enzymes such as the poly-ADP ribose polymerases (PARPs) and the sirtuins that carry out DNA repair and maintain epigenetic homeostasis. Levels of this essential cofactor decline with age in somatic tissues (12), and reversing this decline through the administration of metabolic precursors for NAD^+^ has gained attention as a treatment for disease and strategy for maintaining late life health (13, 14). In this study, we sought to investigate the role of NAD^+^ in the age-dependent decline in fertility and to delineate the role of NAD^+^ and the NAD^+^ consuming enzyme SIRT2 in oocyte quality, post-fertilisation embryo development and fertility.

## Results

### Pharmacological and genetic elevation of NAD^+^ enhances oocyte quality, embryo development, and litter size

In somatic tissues, levels of the metabolite NAD^+^ decline with age (10, 12, 15), and this decline is thought to drive some aspects of physiological dysregulation during ageing. Here, we sought to determine whether this metabolite similarly declined in reproductive tissue with age, and whether this contributed to infertility and declining oocyte integrity, given that these are the oldest cells in the body. To address these questions, we used mice, whose fertility starts to decline around 8 months of age due to oocyte defects that are similar to humans (6). We observed a steep decline in NAD^+^ levels in the ovaries of wild-type mice from the age of 4 months (Fig. 1a, Extended Data Fig. 1), which occurred at a much earlier age than has been described for other tissues (10, 12, 15), consistent with the vulnerability of ovarian function to ageing. We hypothesised that restoring NAD^+^ levels using nicotinamide mononucleotide (NMN) (16), the immediate metabolic precursor to NAD^+^, would raise NAD^+^ in the ovary and restore oocyte quality during ageing. Treatment with NMN by oral gavage raised NAD^+^ levels in the ovaries of 10-month old mice (Fig. 1b), suggesting that this could be one strategy for partially reversing the age-related decline in ovarian NAD^+^.

**Figure 1.**
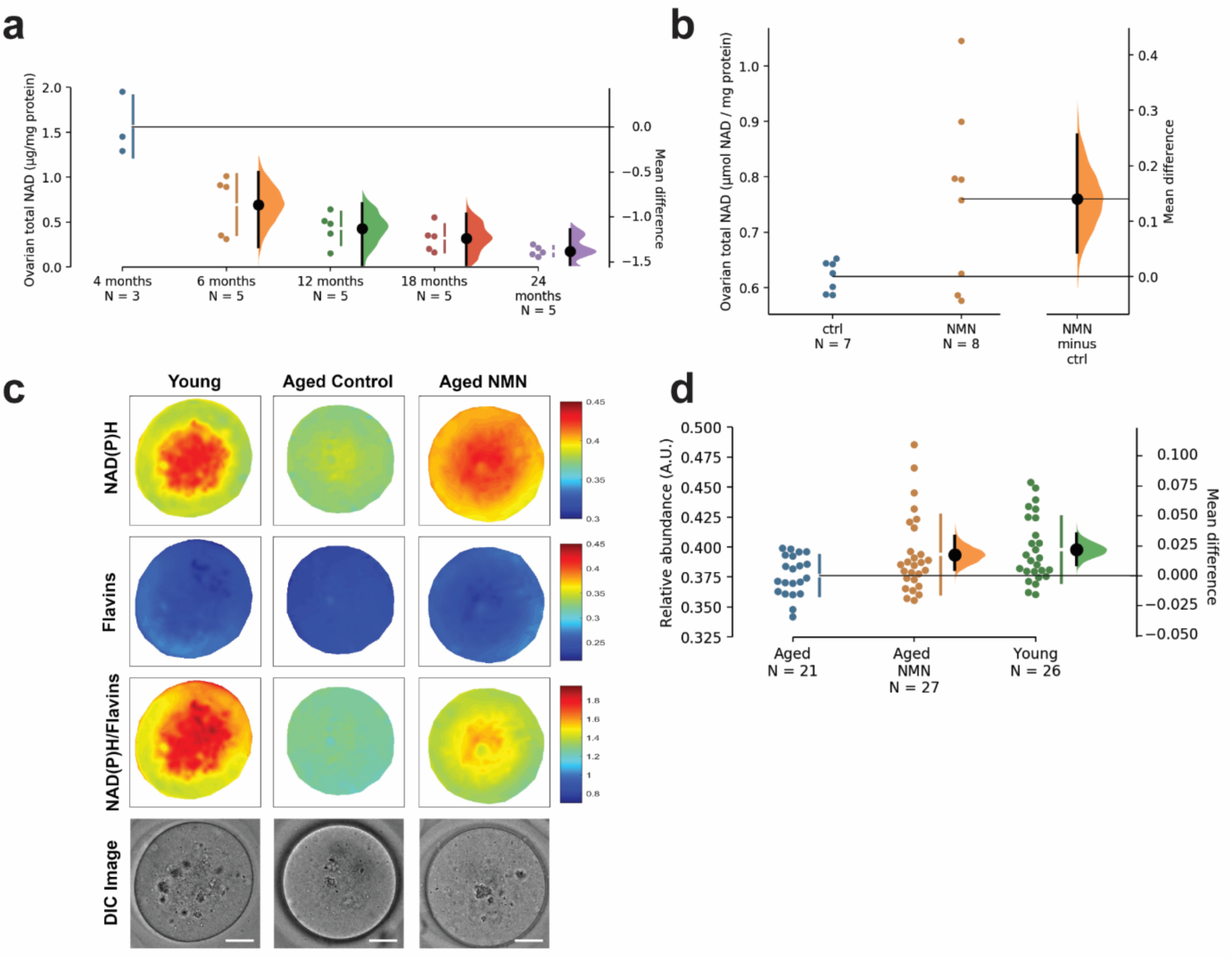
Ovarian and oocyte nicotinamide adenine dinucleotide (NAD^+^) declines with age and can be boosted by oral administration with nicotinamide mononucleotide (NMN). **a**, Ovarian levels of NAD^+^ in the mouse declines with age, which (**b**) can be partially restored in 10 month-old mice following acute treatment (500 mg/kg, 2 hr, oral gavage) with NMN. **c** Multispectral imaging was used to exploit the endogenous fluorescence properties of NADH and NADPH to determine NAD(P)H content in oocytes from young (4-5 weeks old) or aged (12 month old) animals treated with or without NMN through addition to drinking water (2 g/L) for four weeks (scale bar is 20 microns), quantified in **d**. Data throughout this manuscript are presented as modified Cumming (**a, d**) and Gardner-Altman (**b**) plots. Raw data points are shown on the left; the mean difference from control is plotted on a separate axis on the right as a bias corrected and accelerated bootstrap sampling distribution, where 5,000 bootstrap samples are taken, and the confidence interval is bias-corrected. Mean differences are depicted as a dot; the 95% confidence interval is indicated by the ends of the vertical error bar. Data are also presented using traditional null-hypothesis statistics testing in Extended Data.

In addition to changes in NAD^+^ levels in the whole ovary, we next sought to determine whether there was an age-related decline in NAD^+^ levels from individual oocytes, and whether oral NMN treatment could reverse this decline. Accurately measuring NAD^+^ levels in individual oocytes presents analytical challenges due to small sample size, and so we utilised hyperspectral microscopy imaging techniques that exploit the natural fluorescence of NADH and NADPH of individual oocytes. Hyperspectral imaging of autofluorescence allows the characterisation of the most abundant cellular fluorophores, including NADH and flavins (17). The former cannot be spectrally distinguished from NADPH and both are collectively referred to as NAD(P)H, however the autofluorescence of intracellular NADH levels represents the majority of the NAD(P)H signal (18). We treated twelve-month old females with NMN in drinking water (2 g/L) for 4 weeks. At the end of treatment, mature MII oocytes were recovered from the reproductive tracts of females treated with pregnant mare’s serum gonadotropin (PMSG) and human chorionic gonadotropin (hCG). Individual oocytes were then subjected to multispectral microscopy imaging of autofluorescence, followed by unsupervised spectral unmixing to determine the relative abundances of the key native fluorophores, NAD(P)H and FAD, as well as their ratios (Fig. 1c). Consistent with our hypothesis, NAD(P)H levels declined in oocytes from aged (12-month-old) animals, compared to young (4-5-week old) animals, and oral delivery of NMN increased NAD(P)H levels in oocytes from aged animals (Fig. 1d).

To test whether addressing the age-related decline in NAD^+^ would alter oocyte integrity, we treated 14-month old females with NMN via addition to their drinking water (2 g/L) for 4 weeks. To assess the effects of this intervention on oocyte integrity, GV stage oocytes were collected from the ovaries of these animals following stimulation with PMSG, matured *in vitro* to MII, and immunostained to assess spindle structure and chromosome alignment. NMN treatment notably rescued spindle assembly (Fig. 2a), and oocyte yield in aged animals following ovarian hyperstimulation in two strains (C57BL6 and Swiss albino) of mice (Fig. 2b, c). Obesity is also a physiological challenge that results in reduced NAD^+^ (16) and infertility (19), and in obese animals maintained on a high fat diet (HFD) for 5-6 months (Extended Data Fig. 3), NMN also increased oocyte yield (Fig. 2d; Extended Data Fig. 2d).

**Figure 2.**
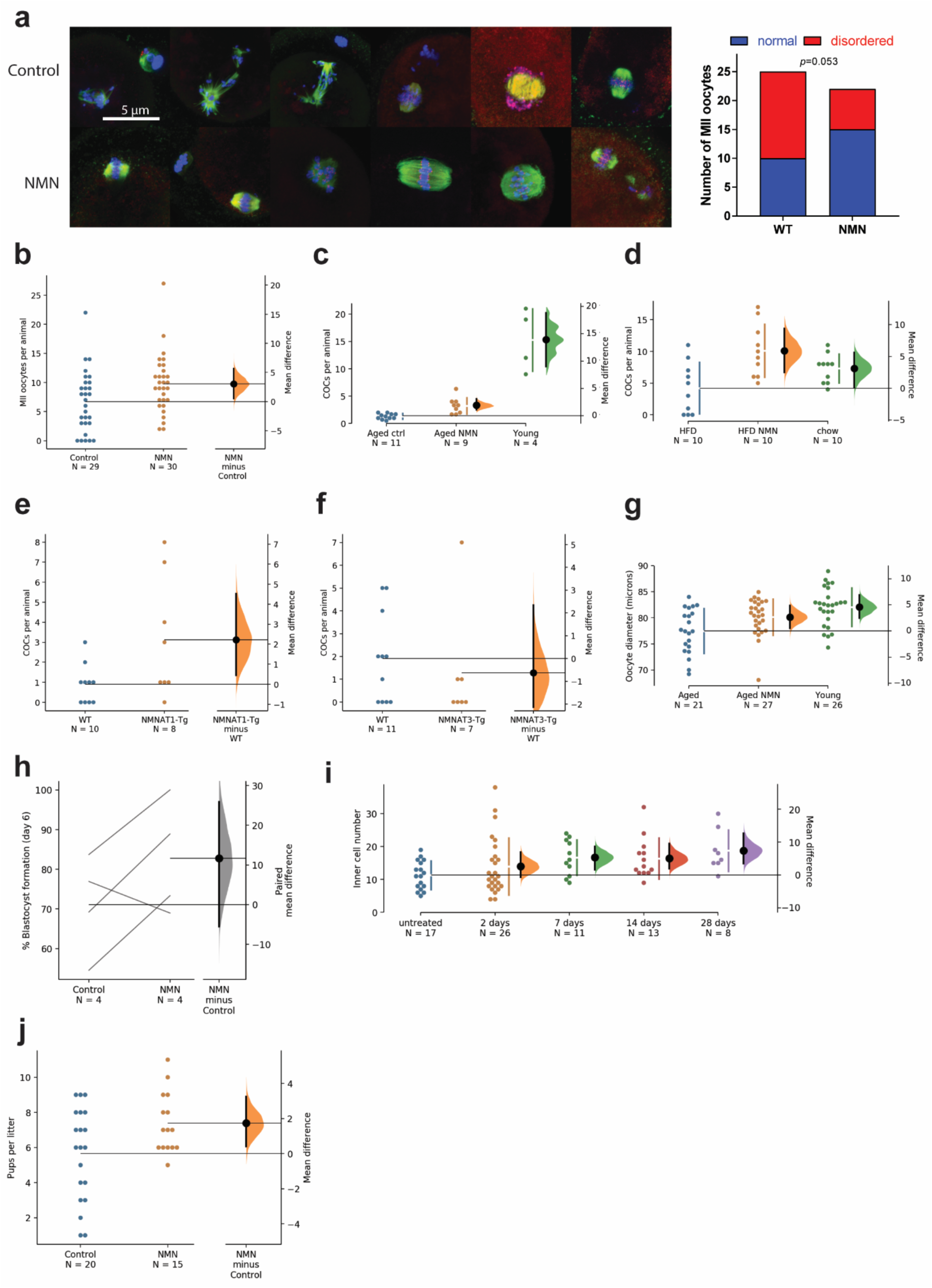
NMN treatments restores oocyte integrity, follicle dynamics, embryo development and pregnancy outcomes. **a**, Chronic treatment (drinking water, 2 g/L, 4 weeks) with NMN from the age of 14 months restores spindle assembly in immunostained oocytes (β-tubulin in green, Hoescht for DNA in blue, kinetochores (ACA) in red, *p*=0.0503 by χ^2^ test). Treatment with NMN improves MII oocyte yield following PMSG and hCG stimulation in (**b**) aged C57BL6 mice, and GV stage oocyte yield in (**c**) 14-16 month old Swiss albino mice, and (**d**) high fat diet (HFD) fed Swiss albino mice after 5-6 months of feeding. **e**, Aged (12-14 month) transgenic mice over-expressing the nuclear NAD^+^ biosynthetic enzyme NMNAT1 have increased oocyte yield, in comparison to (**f**) aged (12-14 month) transgenics overexpressing the mitochondrial NAD^+^ biosynthetic enzyme NMNAT3. **g**. Following 4 weeks of NMN treatment (2 g/L, drinking water) in aged (12-month old) or young (4-6 week old) C57BL6 females, MII oocytes were collected following PMSG and hCG stimulation and oocyte diameter assessed. In a separate cohort, **h** these mature oocytes were used for IVF, with comparisons of percentage blastocyst formation at day 6 of embryo development shown as a slopegraph with each line representing a separate experimental cohort. **i**. 12 month-old C57BL6 females were treated for the indicated times with NMN in drinking water (2 g/L), and MII oocytes collected following PMSG and hCG stimulation and subjected to IVF. At day 6, embryos were fixed and differentially stained to assess inner cell mass. **j**. C57BL6 females (n=10 per group) were treated with NMN (drinking water, 2 g/L) from the age of 10 weeks, and then subjected to timed breeding at the age of 18 weeks, for 5 rounds with 7-8 weeks between rounds until the age of 50 weeks, and the number of pups born per litter recorded.

To further test the importance of NAD^+^ biosynthesis, we studied transgenic strains of animals which over-express the NAD^+^ biosynthetic enzymes NMNAT1 or NMNAT3 (Fig. 2e, f; Extended Data Fig. 4) (20), which are localised to the nucleus and the mitochondria, respectively (21), at the age of 12-14 months. As with NMN treatment, *Nmnat1*^*Tg/+*^ animals yielded more oocytes than their *Nmnat1*^*+/+*^ wild-type littermates (Fig. 2e). In contrast, there was no change in oocyte yield from *Nmnat3*^*Tg/+*^ animals (Fig. 2f), suggesting that the subcellular localisation of NAD^+^ biosynthesis is important for follicular and oocyte function. Given the subcellular localisation of NMNAT1 to the nucleus (21), this would suggest that nuclear NAD^+^ synthesis is important to oocyte development, however another recent study demonstrated an important role for NMNAT2 in oocytes during ageing (22). To test the requirement for NAD^+^ biosynthesis in maintaining normal oocyte function, we next treated GV stage oocytes with FK866, an inhibitor of the NAD biosynthetic enzyme NAMPT (23), and assessed meiotic progression (Extended Data Figure 5). Both germinal vesicle breakdown (GVBD) and polar body extrusion (PBE) were slowed by FK866 treatment, consistent with the idea that NAD^+^ levels are a key determinant of oocyte function.

Next, we sought to determine whether these oocytes from aged, NMN treated animals would have improved performance in embryo development following IVF. Twelve-month old animals were treated with NMN for 4 weeks (2 g/L, drinking water), MII oocytes were collected from oviducts following PMSG and hCG stimulation. Oocytes from NMN treated, aged (12-month old) animals had a larger diameter, comparable to oocytes from untreated aged animals (Fig. 2g). A separate cohort of oocytes were subjected to IVF, and at day 6, the proportion of embryos that reached blastocyst formation was assessed (Fig. 2h), with a trend towards improved blastocyst formation rates. We next sought to determine whether *in vivo* NMN treatment would alter subsequent inner cell mass development of IVF blastocysts, as inner cell mass size is highly predictive of implantation and pregnancy success (24). Twelve-month old mice were treated with NMN in drinking water for 2, 7, 14 or 28 days, and subjected to PMSG and hCG stimulation to promote oocyte release and maturation. These MII oocytes were collected from the oviduct, and subjected to IVF. At day 6, embryos were fixed, and subjected to differential staining to identify the inner cell mass. The length of NMN treatment in animals correlated with improvements in inner cell mass size (Fig. 2i), providing further evidence that oral treatment with NMN intrinsically enhances oocyte quality. Finally, to confirm that this translated to improved fertility outcomes, we treated a cohort of animals with NMN (drinking water, 2 g/L) from 10 weeks of age, and at 18 weeks of age, introduced a male of proven fertility for timed mating. This was repeated every 7-8 weeks until the age of 50 weeks, during which the number of live pups born per litter was recorded (Fig. 2j). Consistent with previous experiments, NMN treatment increased litter size from these animals. Overall, these data from orthogonal pharmacological and genetic approaches show that increasing NAD^+^ enhances ovulation rate, oocyte quality and fertility in aged female mice.

Given these data suggesting a possible use of an orally delivered NAD^+^ raising therapeutic such as NMN to improve oocyte quality and fertility, it was important to assess whether this treatment would adversely affect the health or development of offspring following maternal NMN exposure. In the experiment shown in Fig. 2j, NMN treatment was maintained in the lead-up to pregnancy, throughout pregnancy and during lactation. A cohort of these offspring were maintained to determine if there was any change in offspring health from maternal NMN exposure, and to further uncover whether these offspring would be susceptible to metabolic vulnerabilities, half of these animals were metabolically challenged with high fat feeding following their weaning. Offspring were examined for changes in body weight, body composition, glucose homeostasis, and depression and anxiety-like behaviours (Fig. 3, Extended Data Fig. 6). There were no changes in any of these parameters from maternal NMN exposure, indicating normal development, with the exception of a small but consistent increase in lean body mass from maternal NMN treatment (Fig. 3d). The reason for this change is unclear, and worthy of later investigation.

**Figure 3.**
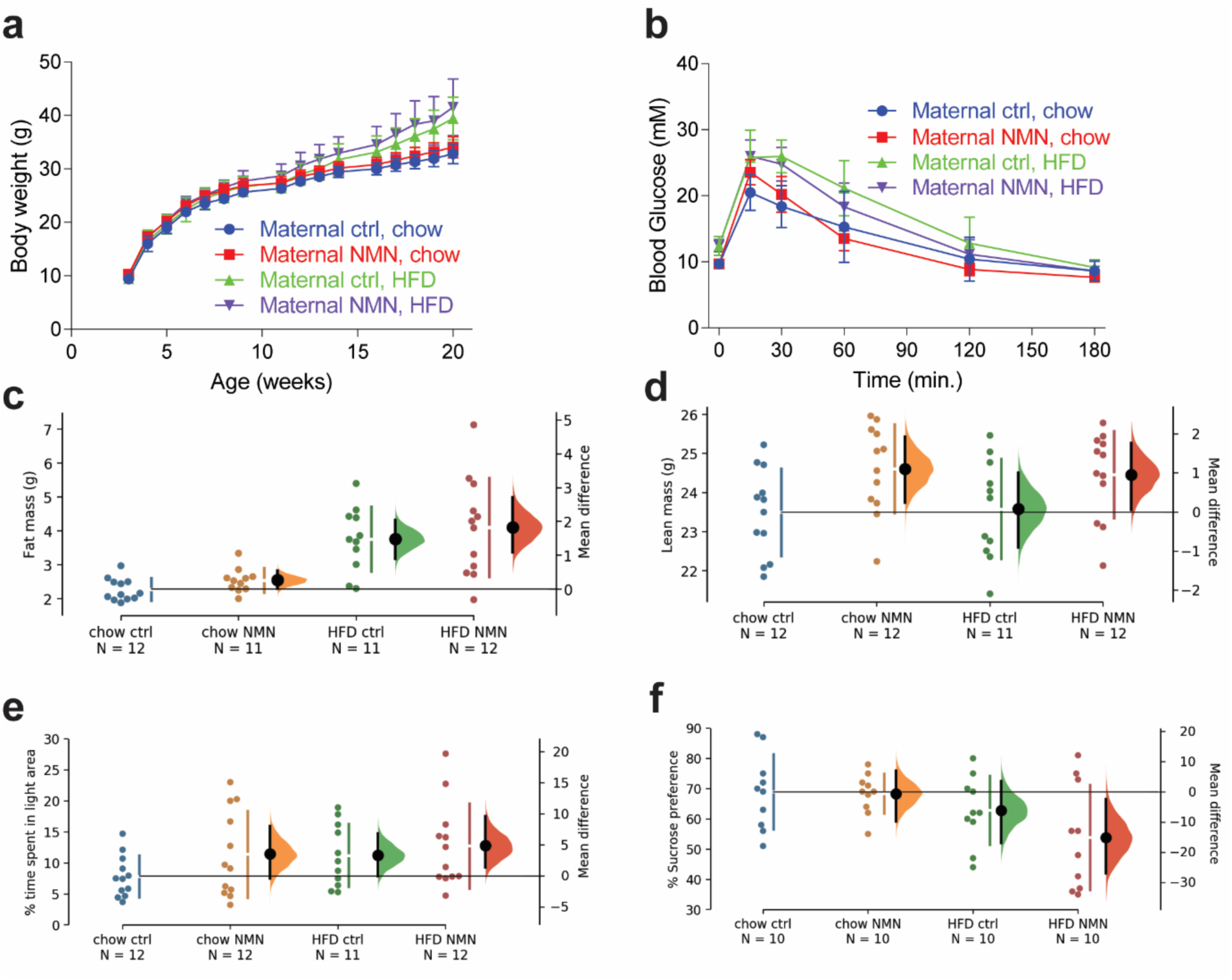
Maternal NMN treatment does not impact growth, metabolism or behaviour in offspring. Male offspring from females treated with or without NMN were maintained on standard chow or high fat diets (HFD). **a**, Body weight was measured on an ongoing basis. **b**, Metabolic homeostasis was measured by glucose tolerance test. **c**, Fat mass and (**d**) lean body mass were assessed by quantitative MRI, with a difference in fat mass from HFD but no effect of maternal NMN treatment, and a small but significant increase in lean body mass with maternal NMN treatment. Behaviour was assessed by the (**e**) light/dark box test for anxiety-like behaviour, and the (**f**) sucrose preference test for depression-like behaviour. Data are also presented using traditional null-hypothesis statistics testing in Extended Data.

### SIRT2 expression in vivo is sufficient but not necessary to recapitulate benefits to oocyte integrity during ageing

Given the ability of NMN to improve aspects of oocyte quality and fertility following *in vivo* treatment, we next sought to determine whether particular enzymes that are critically dependent on NAD^+^ levels for their activity could mediate these benefits. One candidate that we hypothesised for this role is the NAD^+^-dependent deacylase SIRT2. As with all members of the sirtuin family, the activity of this enzyme is critically dependent upon NAD^+^ levels (25), which decline with age (Fig. 1) (12). We were attracted to this candidate due to its previously described role in maintaining processes that are essential to oocytes. We previously showed that the NAD^+^ dependent deacylase SIRT2 stabilises the SAC protein BubR1 (26), which is critical for meiotic progression (27), kinetochore attachment and chromosome segregation in oocytes (28-30). Hypomorphic BubR1 mutants with decreased levels of BubR1 are infertile (11), and levels of BubR1 decline in mouse reproductive tissue (26) and human oocytes with advancing age (31). SIRT2 also maintains genome stability through deacetylation of Cdc20 and Cdh1 required for sustaining the activity of the anaphase-promoting complex (APC) (32), which is an essential oocyte regulator (33, 34). SIRT2 also deacetylates and maintains the activity of the pentose phosphate pathway enzyme glucose-6-phosphate dehydrogenase (G6PD) (35), which protects oocytes against oxidative stress by regenerating the cellular antioxidant glutathione (36).

To test the idea that SIRT2 was involved in or could recapitulate the benefits of NMN treatment, we obtained a strain of *Sirt2*^*Tg/+*^ mice (26) which over-express SIRT2 in all tissues including oocytes (Extended Data Fig. 7a), and assessed oocyte quality at a late reproductive age of 14 months. As with our previous experiments using *in vivo* NMN treatment in aged animals, oocytes were immunostained to assess spindle structure and chromosome alignment. As expected, over 70% of oocytes from reproductively aged wild type (*Sirt2*^*+/+*^) animals had strikingly disordered spindles and poorly aligned chromosomes (Fig. 4a, b), whereas 80% of oocytes from SIRT2 transgenic (*Sirt2*^*Tg/+*^) littermates exhibited normal barrel-shaped bipolar spindles and well-aligned chromosomes. Following hyperstimulation with PMSG, twice as many fully-grown cumulus-enclosed oocytes were obtained from *Sirt2*^*Tg/+*^ animals compared to *Sirt2*^*+/+*^ littermates (Fig. 4c, Extended Data Fig. 8). Given the importance of spindle integrity and chromosome alignment for chromosome segregation, we next tested whether oocytes from aged *Sirt2*^*Tg/+*^ animals might be less prone to aneuploidy, a pathogenomic feature of poor-quality oocytes from aged females. Predictably, aneuploidy rates increased with ageing, from 15% at 3 months of age, to 43% at 16 months of age in *Sirt2*^*+/+*^ wild type oocytes, whereas in oocytes from aged *Sirt2*^*Tg/+*^ littermates the incidence was 20%, comparable to young females (Fig. 4d).

**Figure 4.**
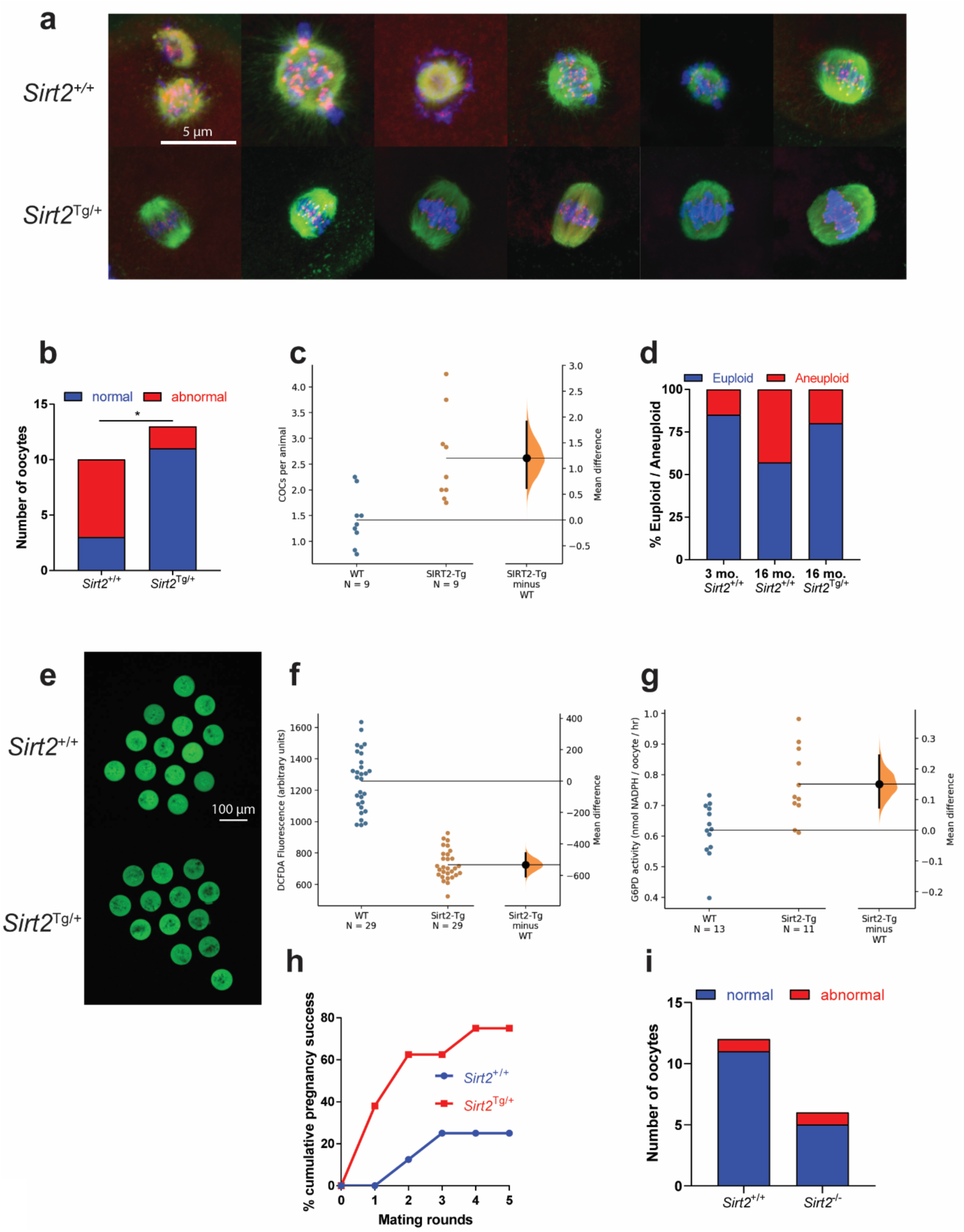
SIRT2 overexpressing transgenic mice have improved oocyte quality. GV stage oocytes were recovered from 14 month-old Sirt2^Tg/+^ C57BL6 mice and matured *in vitro* to the MII stage, following which they were subjected to (**a**) Immunostaining for spindle assembly (β-tubulin in green, Hoescht for DNA in blue, kinetochores (ACA) in red), with (**b**) a quantitative improvement in spindle assembly in Sirt2^Tg/+^ transgenics compared to Sirt2^+/+^ littermates (n=10-13 oocytes per group). **c**, Oocyte yield from reproductively aged (14 month old) PMSG stimulated *Sirt2*^*Tg/+*^ and wild-type *Sirt2*^*+/+*^ littermates (**p=0.0030, n=9 animals per group). **d**, Aneuploidy rates in oocytes from young (2 month old) and aged (16 month old) *Sirt2*^*Tg/+*^ and wild-type *Sirt2*^*+/+*^ littermates (n=30 young wild-type, 7 Sirt2^+/+^ and 5 Sirt2^Tg/+^ oocytes from 4 animals per group). Oocytes from Sirt2^Tg/+^ mice had decreased ROS levels as determined by (**e**) H_2_DCFDA staining, quantified in (**f**), due in part to (**g**) increased G6PD enzyme activity (each data point represents 5 pooled oocytes from 4 animals per group). **h**, Timed mating trials starting from 15 months of age to determine cumulative pregnancy rates, as determined by ultrasound imaging of a foetal heartbeat (p=0.1319 after 5 mating rounds, n=8 animals per group). SIRT2 is sufficient, but necessary for oocyte integrity as (**i**) oocytes from 5 to 6-month old *Sirt2*^*-/-*^ knockout animals display normal spindle assembly when as assessed as in (**a**). Data analysed by two-sided Fisher’s exact test in (**b**) and (**h**). Full statistical analyses in Extended Data Table 1 and Supplementary Information.

Oxidative stress is considered a key driver of oocyte ageing and female infertility (37). SIRT2 deacetylates and maintains the activity of the pentose phosphate pathway enzyme glucose-6-phosphate dehydrogenase (G6PD) (35), which regenerates the antioxidant glutathione through its production of NADPH. Compared to wild type oocytes, *Sirt2*^*Tg/+*^ oocytes from aged animals displayed markedly reduced levels of ROS as determined by staining with the ROS-sensitive fluorescent dye H_2_DCFDA under both young (5-6 month old) unchallenged (Fig. 4e, f) and H_2_O_2_ challenged conditions (Extended Data Fig. 8e, f). Consistent with our hypothesis and these data, we observed increased G6PD enzyme activity in these oocytes (Fig. 4g).

Given the improved characteristics of oocytes from aged *Sirt2*^*Tg/+*^ animals, we next sought to determine whether this translated into improved fertility. Animals from this strain were aged to 15 months, well past the normal end of fertility for this strain of around 8 months, and subjected to timed breeding trials to determine pregnancy rates, as defined by the presence of a fetal heartbeat. Consistent with very low fertility at this age, only 25% of wild type *Sirt2*^*+/+*^ females achieved a pregnancy over 5 mating rounds. Notably, in *Sirt2*^*Tg/+*^ females pregnancy rates tripled to 75% (Fig. 4h). We were unable to ascertain litter size and live birth rates in this study, due to high rates of rapid parental infanticide observed in these aged animals, however increased pregnancy rates are indicative of improved fertility during ageing. Taken together, these data demonstrate that the NAD^+^ dependent deacylase SIRT2 is sufficient to maintain ovarian function and female fertility during ageing.

Although these data suggested that SIRT2 was sufficient to recapitulate the benefits of NMN to fertility, we next sought to determine whether SIRT2 was also obligatory for maintaining normal oocyte function, through the use of *Sirt2*^−/−^ knockout animals. Oocytes from whole body *Sirt2*^−/−^ knockout mice at the age of 5-6 months displayed completely normal spindle assembly and maturation (Fig. 4i), indicating that at a younger age where NAD is replete, SIRT2 is not essential for accurate spindle assembly, or that there is redundancy in the role of SIRT2 with other yet to be identified factors. These *in vivo* results from *Sirt2* knockout animals are in contrast to studies of *in vitro* morpholino knockdown of *Sirt2* (28) or chemical inhibition of SIRT2 in oocytes (38), where depletion or inhibition results in spindle assembly defects and aneuploidy (28, 29). The discrepancy between these results may be due to the compensatory, upregulation of other factors in constitutive knockout animals, versus the acute depletion of SIRT2 in oocytes during *in vitro* maturation. Together, these data suggest that SIRT2 is sufficient, but not required to improve oocyte quality during ageing.

### In vitro treatment with NMN enhances embryo development during ageing or in a minimal growth environment

These data show that tissue NAD^+^ levels are critical for oocytes and female fertility during ageing. Next, we asked whether elevating NAD^+^ might also benefit pre-implantation embryo development. If so, *in vitro* treatment with NAD^+^ precursors might enhance embryonic developmental milestones and benefit IVF outcomes for reproductively aged females, enabling its rapid clinical translation. To test this, IVF was performed using *in vivo* matured oocytes from reproductively aged (12 month) or young (4 weeks) females. Following IVF, embryos were cultured in normal, replete embryo culture medium in the presence or absence of NMN. Consistent with the idea that a deficiency in NAD^+^ levels with increasing age drives poor reproductive outcomes, supplementation of medium with NMN improved blastocyst formation in embryos derived from oocytes from aged, 12 month old females (Fig. 5a, Extended Data Fig. 9, 10), but not in embryos arising from oocytes from young, 4-week old females (Extended Data Fig. 10) which naturally exhibit high developmental competence. To further assess whether NMN could rescue embryo development under challenged conditions, embryos from young animals were maintained in simple culture media, which accentuates culture stress and restricts embryo development (39). Consistent with results using oocytes from aged animals, the addition of NMN to simple media improved blastocyst cell number, an indicator of implantation success (Fig. 5b).

**Figure 5.**
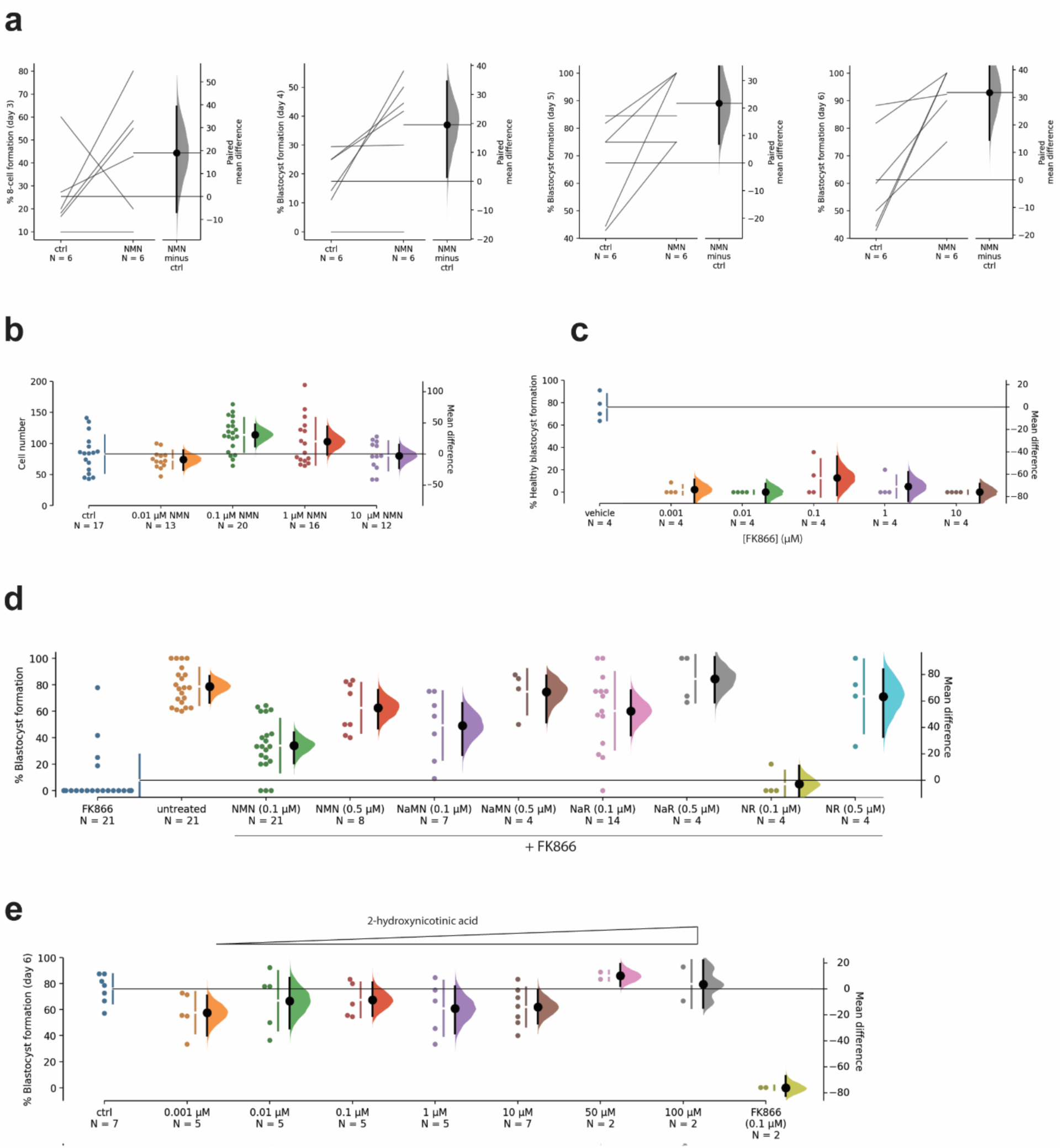
*In vitro* NMN treatment enhances embryo formation under aged or challenged conditions. MII oocytes from (**a**) aged (12 month old) mice were subjected to IVF, and post-fertilisation embryos maintained in media containing 1 μM NMN until day 6 of embryo development, with the percentage of embryos reaching developmental milestones (8-cell or blastocyst) shown for days 3-6. The addition of NMN could also enhance cell count at 92 hr in blastocysts from young animals matured in (**b**) simple defined media, which induces culture stress. **c**, Treatment with the NAMPT inhibitor FK866 at the indicated concentrations causes embryo death at day 6 (% surviving blastocysts shown), which (**d**) can be rescued by treatment with the NAD^+^ precursors NMN, nicotinic acid mononucleotide (NaMN), nicotinic acid riboside (NaR) and nicotinamide riboside (NR). **e**, Treatment with the NaPRT inhibitor 2-hydroxynicotinic acid has a minimal effect on blastocyst formation, compared to FK866 treatment (day 6 data shown).

Mammalian cells have the capacity to generate NAD^+^ via several biosynthetic routes and intermediate precursors, including from nicotinamide in the recycling pathway, nicotinic acid in the Preiss-Handler pathway (40, 41), tryptophan and quinolinic acid in the *de novo* pathway (42, 43), and nicotinamide riboside in the nicotinamide riboside kinase (NRK) pathway (44). NMN is an intermediate in the recycling pathway, where the enzyme nicotinamide phosphoribosyltransferase (NAMPT) generates NMN from nicotinamide, and it is also an intermediate in the NRK pathway, where the enzyme NRK phosphorylates nicotinamide riboside (NR) into NMN (44). To assess whether NAD^+^ synthesis via endogenous NMN production in the recycling pathway plays a role in embryo development, we cultured embryos in the presence of FK866, an inhibitor of the enzyme NAMPT (23). FK866 treatment induced blastocyst degeneration on day 6 of culture (Fig. 5c), and this could be rescued by NMN co-treatment (Fig. 5d). Interestingly, the blastocyst degeneration induced by FK866 treatment could also be partially or completely rescued through treatment with other NAD^+^ precursors including nicotinic acid mononucleotide (NaMN), nicotinic acid riboside (NaR) and nicotinamide riboside (NR) (Fig. 5d). These data suggest that while NAD^+^ production requires NMN via NAMPT in the recycling pathway in the developing embryo, the NRK and Preiss-Handler pathways are also present and capable of compensating for this deficiency. In contrast to the embryo degeneration imposed by NAMPT inhibition, treatment with 2-hydroxynicotinic acid, an inhibitor of the Priess Handler pathway enzyme nicotinic acid phosphoribosyltransferase (NaPRT) (45), did not affect embryo development (Fig. 5e). Together, these data suggest that the Preiss-Handler pathway is present and capable of compensating for NAMPT inhibition, but is not required for NAD^+^ biosynthesis in embryos under normal circumstances.

NAD^+^ plays a prominent role as a redox cofactor used in fundamental metabolic reactions required to sustain life. It is also consumed as a substrate by several enzymes which have been described to play a role in biological ageing, most notably the sirtuins, which are thought to be critically dependent on NAD^+^ availability. To determine the contribution of this class of enzymes to embryo development and the effects of NMN, we next grew embryos in the presence of the small molecule sirtuin inhibitors sirtinol (46) and splitomicin (47) under simple defined growth media conditions which can induce culture stress (39). Both inhibitors reduced both blastocyst formation and cell numbers (Fig. 6a-b, Extended Data Fig. 12). We next sought to determine whether the effects of NMN were dependent on the activity of the sirtuins through co-treating embryos under simple defined media growth conditions (39) with NMN and sirtinol (Fig. 6c). The decline in embryo cell number induced by sirtinol treatment was partially reversed by NMN co-treatment (Fig. 6c), suggesting that the ability of NMN to enhance cell number and developmental milestones was not solely dependent upon the targets of this inhibitor. While these data suggest that the activity of the sirtuins are essential to embryo development, they do not suggest that they are the primary target of NMN in enhancing post-fertilisation embryogenesis.

**Figure 6.**
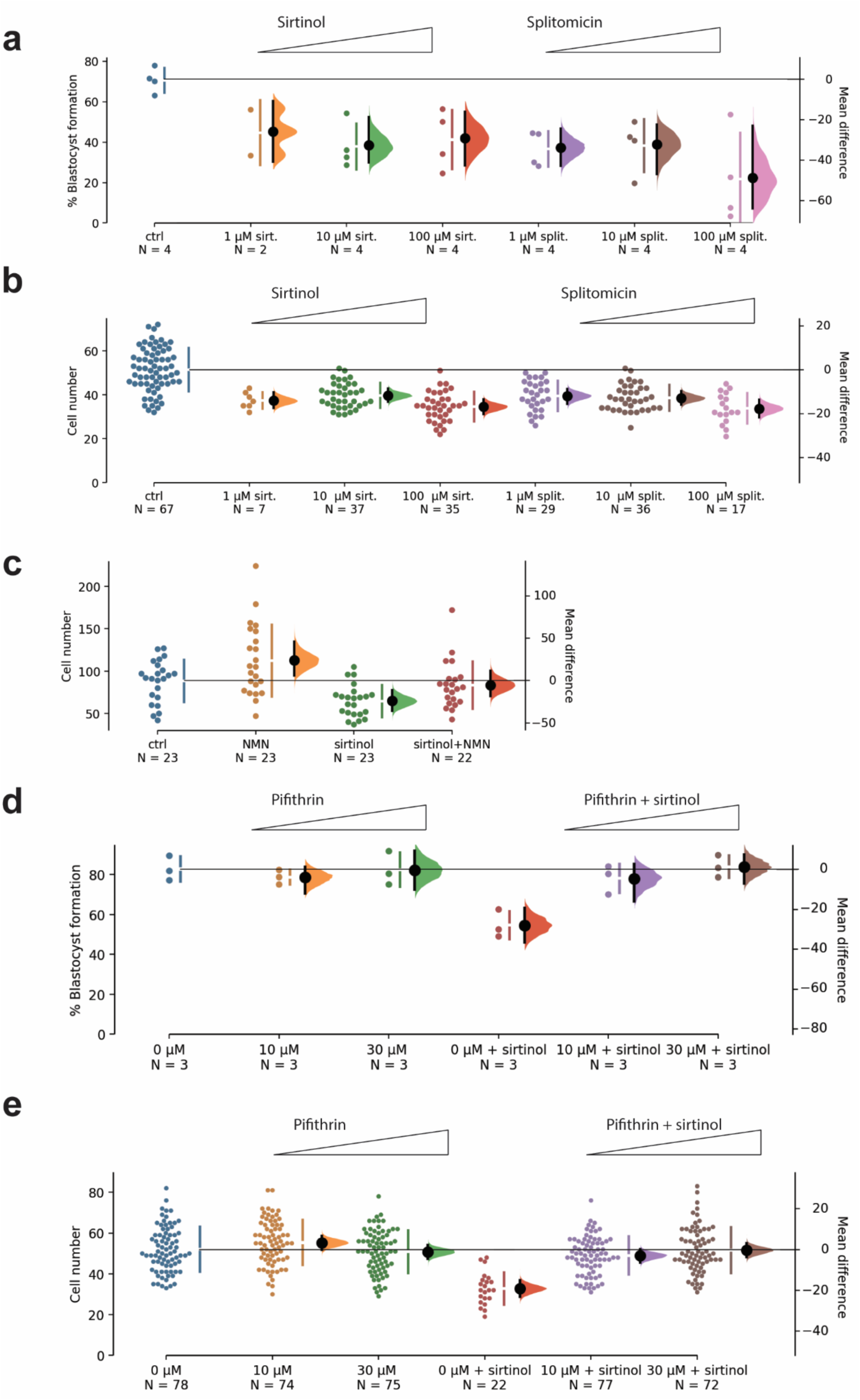
Benefits of NMN to blastocyst formation are independent of sirtuin activity, which is required for p53 dependent embryo formation. Treatment with the small molecule sirtuins inhibitors sirtinol or splitomicin (**a**) inhibits blastocyst formation, with (**b**) decreased cell count in those blastocysts that are formed. **c**, Co-treatment of sirtinol treated embryos with NMN rescues this reduction in cell count, indicating that the benefits of NMN are partially independent of sirtuins activity. Treatment with the p53 inhibitor pifithrin rescues (**d**) blastocyst formation and (**e**) cell count in embryos treated with sirtinol. Data from embryos maintained in defined simple media, fixed at 92 hr post-fertilisation.

Given the ability of sirtuin inhibitors to reduce blastocyst quality, we next sought to determine the pathway through which inhibition of sirtuins would restrict embryo growth. The most prominently studied member of the sirtuins is SIRT1, which has deacetylase activity towards p53 to inhibit its activity and prevent apoptosis (48, 49). p53 activity is increased in embryos produced by IVF compared to *in vivo* derived embryos, likely due to culture stress, and its heterozygous or homozygous genetic deletion overcomes the effects of culture stress on blastocyst development (39, 50). Activation of p53 is detrimental as it specifically inhibits proliferation and induces apoptosis of the inner cell mass, which goes on to form the developing fetus (51-53). Here, we observed that treatment with the p53 inhibitor pifithrin (54) overcame sirtinol-mediated reduction in blastocyst development (Fig. 6d) and cell number (Fig. 6e), suggesting that p53 dependent hypotrophy is sirtuin mediated, a result that is consistent with previous findings on the mechanism of apoptosis in other cell types by sirtinol (55).

## Discussion

There is an ongoing trend across the developed world to defer pregnancy until later in life. Combined with the steady decrease in female fertility beyond the middle of the third decade of life, this has led to falling fertility rates and a steady increase in demand for assisted reproduction technologies. Despite maternal age being the greatest clinical challenge for reproductive medicine, the mechanisms through which oocyte quality decline as women age remain largely unclear and there are no therapeutic treatments. In the current study, we provide evidence for NAD^+^ availability as an important determinant of fertility in aged females. We show that levels of NAD^+^ rapidly decline in the ovary with age, and our data demonstrate that NAD^+^ repletion using the NAD^+^ precursor NMN in mice enhances ovulation rate, reduces oocyte spindle defects, improves oocyte developmental competence *in vivo* and embryo development during IVF, culminating in improved fertility.

The present study is also the first to implicate the NAD^+^ pathway in regulating ovulation rate, which applies to a poly-ovulatory species such as the mouse. While we measured NAD^+^ levels in the ovary in this study, we do not exclude the possibility that NMN exerts benefits especially to folliculogenesis through direct interactions with tissues other than the ovary, as we delivered NMN through systemic dosing. NMN may exert primary effects directly in the ovary, or it may instead be possible that systemic improvements in vascular function (56), metabolic homeostasis (13, 16), mitochondrial function (10) and stem cell function (57) or signalling in the brain could lead to secondary improvements in ovarian function. These findings should especially be viewed in light of the well-studied role of NAD^+^ raising molecules in maintaining late-life health (14) and the biology of ageing. Regardless, these results provide support for the premise that age-related reductions in NAD^+^ availability contribute to female infertility, and that improved oocyte integrity and/or oocyte yield following superovulation following pharmacological restoration of NAD^+^ through the administration of metabolic precursors opens a therapeutic window for the treatment of age-related infertility, and improving the clinical success rates of IVF.

While NAD^+^ is a prominent molecule used as a cofactor or substrate across a range of reactions, we sought to address the hypothesis that the benefits of treating with NMN to oocyte integrity might in part be mediated by the NAD^+^ dependent deacylase SIRT2. The benefits of *in vivo* NMN treatment could largely be recapitulated by transgenic overexpression of *Sirt2* in aged animals, although its constitutive deletion had no adverse impact on oocyte integrity, at least in the younger animals studied here. SIRT2 plays a role in the maintenance of microtubule-kinetochore attachments through its deacylation and stabilisation of BubR1 (26, 29), a process that ensures fidelity in chromosome separation through ensuring the bipolar orientation of chromosomes to the spindle. Consequently, in oocytes from *Sirt2*^*Tg/+*^ overexpressing animals, increases in kinetochore – microtubule stability likely contributed to augmented chromosome alignment and improved fertility. These observations are corroborated by separate *in vitro* studies where morpholino-mediated *Sirt2* knockdown or chemical inhibition of SIRT2 in oocytes resulted in severe spindle defects and chromosome disorganization (29, 38). These results are supported by the apparent lower rates of aneuploidy in oocytes from aged *Sirt2*^*Tg/+*^ animals in the present study. Surprisingly and in contrast to studies of *in vitro* knockdown of *Sirt2* following morpholino microinjection into oocytes, we observed that oocytes from constitutive *Sirt2* knockout mice maintained normal spindle assembly and chromosome organisation, suggesting the existence of overlapping mechanisms for this process that may compensate during development, and that SIRT2 is sufficient but not necessary for spindle assembly, at least at a younger age where NAD^+^ is replete.

Having demonstrated that *in vivo* NMN treatment increased ovulation rate, improved oocyte quality and overall litter size, we assessed the effect of supplementing embryo culture media with NMN. The present study established that culturing embryos from aged female mice in the presence of NMN improved blastocyst formation. Similarly, NMN increased blastocyst cell number in embryos from old animals cultured in replete embryo medium and in embryos from young animals grown in nutritionally suboptimal medium (39). Importantly, NMN did not confer any benefit to embryos from young animals maintained under standard embryo culture conditions, supporting the idea that this intervention addresses an age-related deficit in NAD^+^ levels. These findings are highly relevant to the clinical practice of IVF. In addition to age-related issues of decreased oocyte numbers and oocyte integrity; mitotic aneuploidy (58) and poor preimplantation embryo development (59) limits the number of euploid blastocysts available for transfer. Increased mtDNA copy number (60) and reduced gene expression regulating cell cycle control (61) relative to younger patients have been implicated in reduced viability in embryos derived from females of advanced maternal age. This manifests in reproductively aged females having notably fewer viable euploid blastocysts for transfer, culminating in IVF live birth rates falling from 19.2% in women under 30, 12.4% in 35-39, and 4.6% in women 40-44 years of age (62). These Australian and New Zealand data underscore the significant clinical demand for a therapeutic that can assist in improving pregnancy success rates in reproductively aged women.

In this work we also interrogated the contributions of the different NAD^+^ biosynthetic pathways present in mammals to embryo health. Our experiments focused on NMN, an intermediate in the NAD^+^ recycling pathway produced by the enzyme NAMPT, and also the product of the NRK enzymes, which feed into the same pathway. Running separately to the recycling or NAD^+^ pathways, the Priess-Handler pathway and *de novo* pathway both utilise the intermediate NaMN via the enzymes NaPRT and QPRT, respectively. We found that blastocysts underwent degeneration when exposed to the NAMPT inhibitor FK866 (23), but not the NaPRT inhibitor 2-hydroxynicotinic acid (45). NMN co-treatment overcame the block on NAMPT by FK866, however the intermediates NR, NaR and NaMN also overcame the effects of FK866. This suggests that the NRK pathway, which utilises NR and NaR (44, 63), and the Priess-Handler and *de novo* pathways which utilise NaMN (40, 41) are present in the developing embryo, but their activity is not essential to normal embryo growth. The results also shed light on the opportunity to enhance oocyte and embryo health using multiple NAD^+^ boosting compounds.

NAD^+^ raising compounds are frequently studied in terms of their ability to improve the activity of the NAD^+^ dependent sirtuin class of enzymes. We found that two structurally unrelated small molecule inhibitors of the sirtuins drastically reduced embryo development. Given that sirtuin expression decreases as embryo development progresses (53) and NMN co-treatment could partially reverse this block suggests that the effects of NMN may be mediated only in part through this class of enzymes. It is likely that increasing NAD^+^ through NMN treatment enhances embryo development through many processes, including maintaining redox homeostasis, electron transport and oxidative phosphorylation in the mitochondria, nutrient catabolism, facilitation of DNA repair, and as we show here, maintaining activity of the sirtuins. We showed that reductions in embryo development during sirtinol treatment were mediated through p53, which blocks blastocyst development (53) and formation of the inner cell mass (51, 52), particularly during suboptimal culture conditions (50). This mechanism likely reflects the role of SIRT1 as a deacetylase for p53, maintaining it in an inactive state and preventing apoptosis (48).

Together, this work represents a clinically tractable pharmacological intervention to non-invasively treat female infertility caused by a loss of oocyte viability or depletion of ovulation yield in reproductively aged females. These findings have immediate implications for the clinical treatment of infertility. We envisage that this work could lead to the development of an orally delivered therapeutic that improves oocyte quality for improving natural conception or the success of IVF. This would present an opportunity to transcend the need for, or enhance the success of fertility treatments including IVF. In addition, this work could also enhance the success rates of existing IVF protocols, through improving embryo culture conditions to grow embryos to a later stage of development. Providing an intervention at this critical step of IVF would make a clinically relevant difference to IVF success rates in the high proportion of IVF patients who are reproductively aged. Any intervention that would improve the success rates of IVF would also lead to cost savings and lower the emotional stress of failed IVF rounds or subsequent miscarriage which can lead to long term psychological and social issues including depression and relationship breakdown. This would represent the first intervention for enabling women with poor oocyte quality to have children with their own genetic make-up as currently, these women have no alternative but to use donated oocytes. Future studies should aim to test NAD^+^ raising compounds in a clinical setting, both as an oral therapeutic, and as an additive to embryo media during IVF, to test the relevance of these findings to human infertility.

## Methods

Data in this manuscript are presented using modified versions of Gardner-Altman and Cumming estimation plots obtained using the recently developed DABEST data analysis package (64). Raw data points are shown on the left; the mean difference from control is plotted on a separate axis on the right as a bias corrected and accelerated bootstrap sampling distribution (65), where 5,000 bootstrap samples are taken, and the confidence interval is bias-corrected. Mean differences are depicted as a dot; the 95% confidence interval is indicated by the ends of the vertical error bar. Traditional null-hypothesis statistics testing based analyses for each figure are provided in Extended Data.

Detailed methods are available in Supplementary Methods. Further detail on statistical analyses are available in Extended Data Table 1 with all calculations available in Supplementary Information (.xml files).

## Supplementary Information

Supplementary Information 1 provides detailed methods for all experiments, including statistical design. Detailed statistical analysis calculations are provided as Supplementary Information (Data Analysis). Summarised statistics for all figures are also available in Extended Data Table 1.

## Funding and acknowledgments

This work was supported by the National Health and Medical Research Council (NHMRC) of Australia, through grants APP1103689 and APP1122484 to LEW, DAS and HAH, APP1139763 to RBG, LEW and KAW, APP1044295 to MJM and DAS, APP1066172 to NT, DGLC and DAS, and a Career Development Fellowship to LEW (APP1127821). It was also supported by the Australian Research Council (ARC) grants DP170101863 and CE140100003 to EMG. The salary and experimental costs of MB and DMG working in the lab of RBG and LEW was partly supported by Jumpstart Fertility. KS is an employee of Jumpstart Fertility. We gratefully acknowledge assistance from the UNSW Biological Resource Centre. We also wish to thank the Solina Chau foundation and Mr Hejun (Steven) Zhang for their philanthropic support.

## Author contributions

LEW, HAH and DAS conceived of this study and obtained funding. LEW, HAH, DAS, RBG, KAW, CO, MJM and EG designed and supervised experiments, and analysed and interpreted results. WHJH, DRL, MJB, DG, AR, KS, JB, WGNL, ASAW, DR, CL, JM, NY, LL, RMW, LEQ, SC, LJK, SB, XJL, SM, JMC, AH carried out experiments and analysed results. TA generated *Nmnat3* transgenic mice. ASM assisted in statistical analysis. NT, DGLC, provided critical feedback. LEW wrote and prepared this manuscript.

## Author information and disclosures

LEW, HAH and DAS are co-founders, shareholders, directors and advisors of Jumpstart Fertility Inc, which was founded to develop the work described here. The salaries of MJB and DG were paid by contract research from Jumpstart Fertility to UNSW. KS is an employee of Jumsptart Fertility. WHJH and DRL are shareholders of Jumpstart Fertility. LEW and DAS are also advisors and shareholders in EdenRoc Sciences (Metro Biotech NSW, Metro Biotech, Liberty Biosecurity), and in Life Biosciences LLC and its daughter companies (Jumpstart Fertility, Continuum Biosciences, Senolytic Therapeutics, Selphagy, Animal Biosciences). LEW is an advisor and shareholder in Intravital Pty Ltd. DAS is also a founder, equity owner, advisor, director, consultant, investor and/or inventor on patents licensed to Vium, Jupiter Orphan Therapeutics, Cohbar, Galilei Biosciences, Wellomics, EdenRoc Sciences (and affiliates Arc-Bio, Dovetail Genomics, Claret, Revere Biosciences, UpRNA, MetroBiotech, Liberty Biosecurity, Life Biosciences (and affiliates Selphagy, Senolytic Therapeutics, Spotlight Therapeutics, Animal Biosciences, Iduna, Continuum, Jumpstart Fertility). He is an inventor on a patent application filed by Mayo Clinic and Harvard Medical School that has been licensed to Elysium Health. For details see https://genetics.med.harvard.edu/sinclair/

---

**Extended Data Figure 1.**
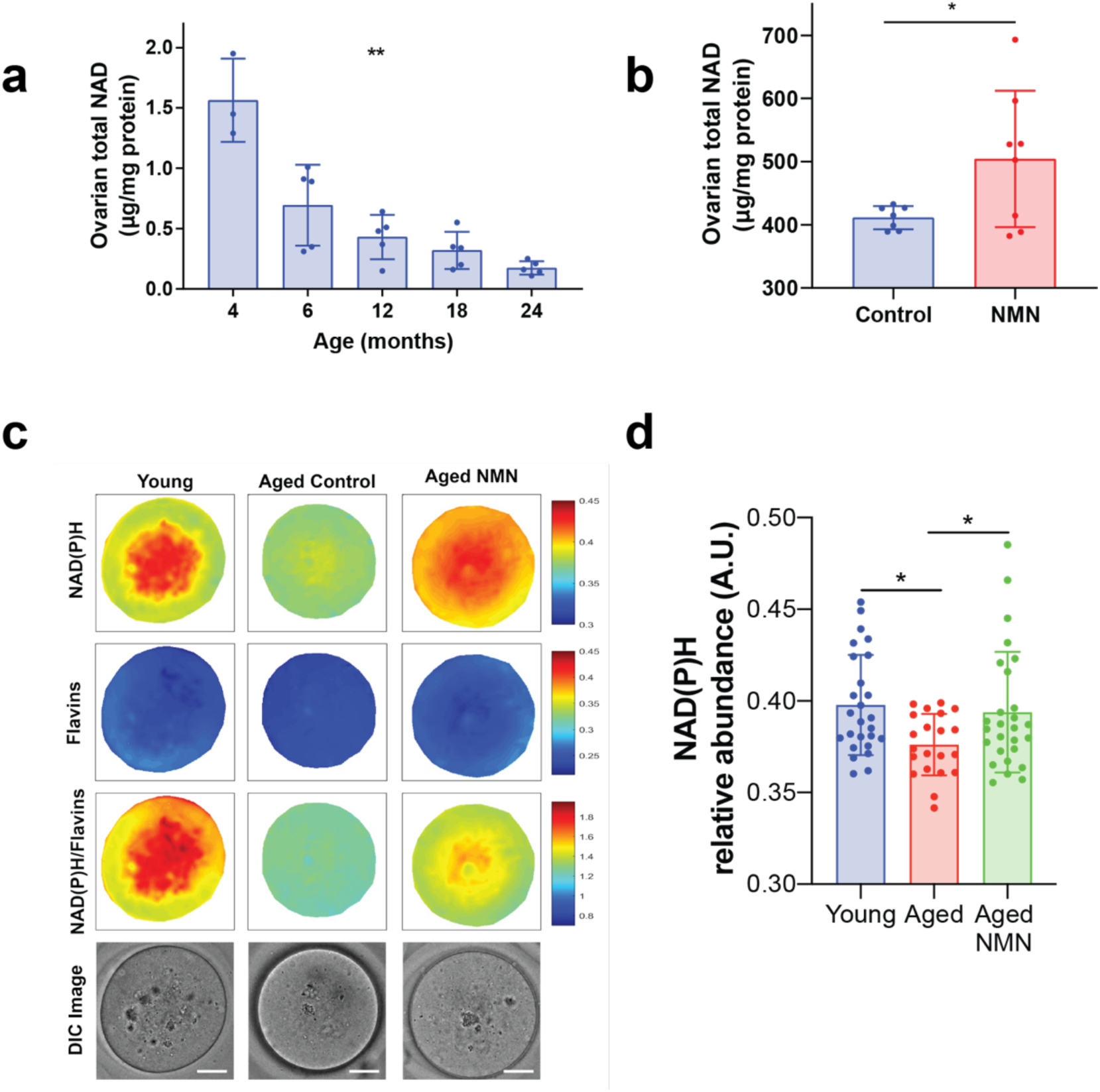
Data from main text Figure 1 presented using traditional null-hypothesis statistics testing. **a**, Ovarian levels of NAD^+^ in the mouse declines with age (\*\**p*=0.0055, *n=*3-5 per age), which (**b**) can be partially restored in 10 month-old mice following acute treatment (500 mg/kg, 2 hr, oral gavage) with NMN (\**p*=0.0433, n=6-7 per group). **c** Multispectral imaging was used to exploit the endogenous fluorescence properties of NADH and NADPH to determine NAD(P)H content in oocytes from young (4-5 weeks old) or aged (12-month old) animals treated with or without NMN through addition to drinking water (2 g/L) for four weeks (scale bar is 20 microns), quantified in **d**. Data analysed by one-way ANOVA in (**a**) and (**d**) with Holm-Sidak post-test in (**d**), analysed by two-way *t*-test in (**b**).

**Extended Data Figure 2.**
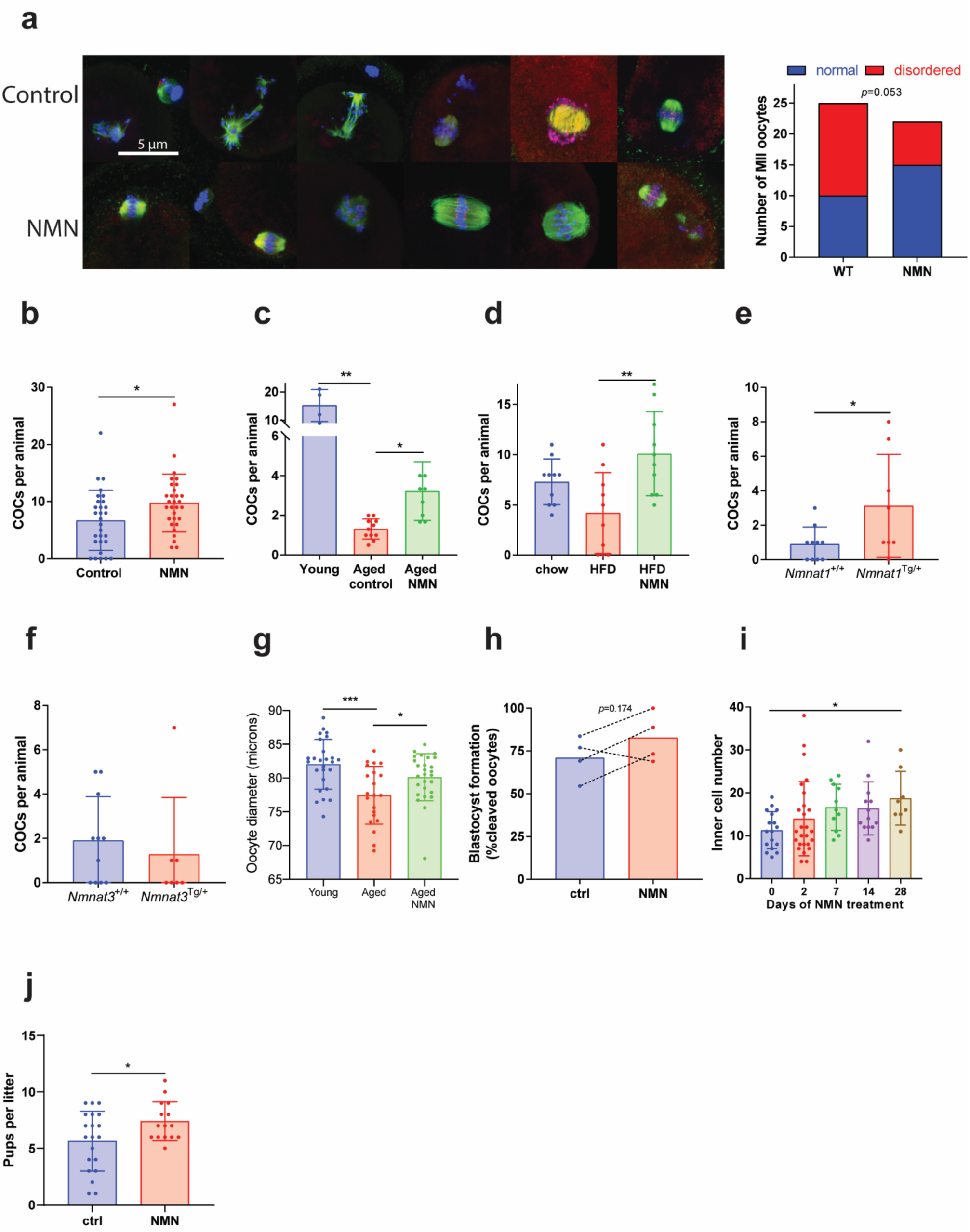
Data from main text Figure 1 presented using traditional null-hypothesis statistics testing. Chronic treatment (drinking water, 2 g/L, 4 weeks) with NMN from the age of 14 months (**a**) restores spindle assembly in immunostained oocytes (β-tubulin in green, Hoescht for DNA in blue, kinetochores (ACA) in red, *p*=0.0503 by χ^2^ test). Treatment with NMN improves oocyte yield following ovarian stimulation in (**b**) aged (12 month old) C57BL6 mice (\**p*=0.0211, *n*=29-30 animals per group), (**c**) 14-16 month old Swiss albino mice (\*\*\**p*=0.0004, \**p*=0.0295 n=4-11 per group), and (**d**) high fat diet (HFD) fed Swiss albino mice after 5-6 months of feeding (\*\**p*=0.0031, *F*=6.746 *n*=10 per group). **e**, Aged (12-14 month) transgenic mice over-expressing the nuclear NAD^+^ biosynthetic enzyme NMNAT1 have increased oocyte yield from ovaries following PMSG stimulation (\**p*=0.0416, *n*=8-10 per group), in comparison to (**f**) aged (12-14 month) transgenics overexpressing the mitochondrial NAD^+^ biosynthetic enzyme NMNAT3. **g**. Following 4 weeks of NMN treatment (2 g/L, drinking water) in aged (12-month old) or young (4-6 week old) C57BL6 females, MII oocytes were collected following PMSG and hCG stimulation and oocyte diameter assessed. In a separate cohort of 12-month old animals, mature oocytes were used for IVF, with **h** comparisons of percentage blastocyst formation at day 6 of embryo development shown as a slopegraph with each line representing a separate experimental cohort. **i**. 12 month-old C57BL6 females were treated for the indicated times with NMN in drinking water (2 g/L), and MII oocytes collected following PMSG and hCG stimulation and subjected to IVF. At day 6, embryos were fixed and differentially stained to assess inner cell mass. Data analysed by Kruskal Wallis test in (**a**) and (**g**); Mann-Whitney U-test in (**e**), one-way ANOVA with Holm-Sidak in (**c**), (**d**), (**g**) and (**i**), unpaired two-tailed t-test in (**f**). Data presented as mean ± s.d; individual data points are shown. Further statistical detail in Extended Data Table 1 and Supplementary Information.

**Extended Data Figure 3.**
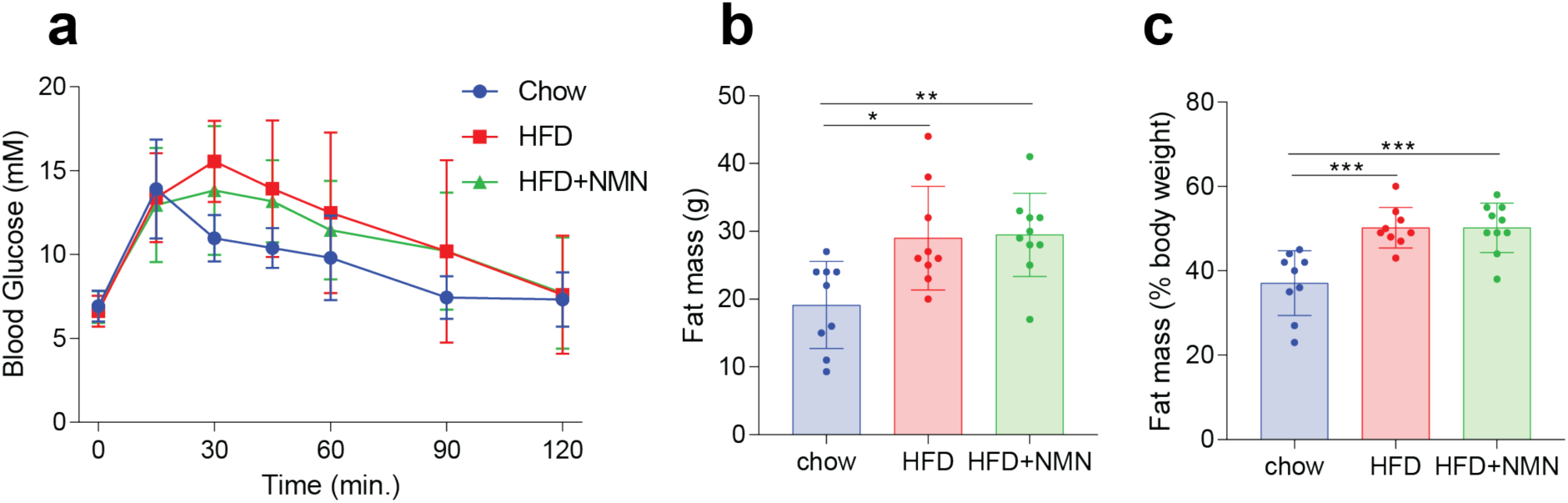
Obesity and metabolic dysfunction in high fat diet (HFD) fed animals. Swiss albino mice were maintained on a standard chow or HFD for 5-6 months in the presence or absence of NMN treatment (2 g/L, drinking water). **a**, Glucose tolerance test (2 g/kg, i.p.), (**b**) quantitative MRI for fat mass, and (**c**) fat mass expressed as a percentage of total body weight. *n*=9-10 animals per group, **p<0.01 one-way ANOVA with Hold-Sidak multiple comparison test. Data are mean ± s.d. Further statistical detail in Extended Data Table 1.

**Extended Data Figure 4.**
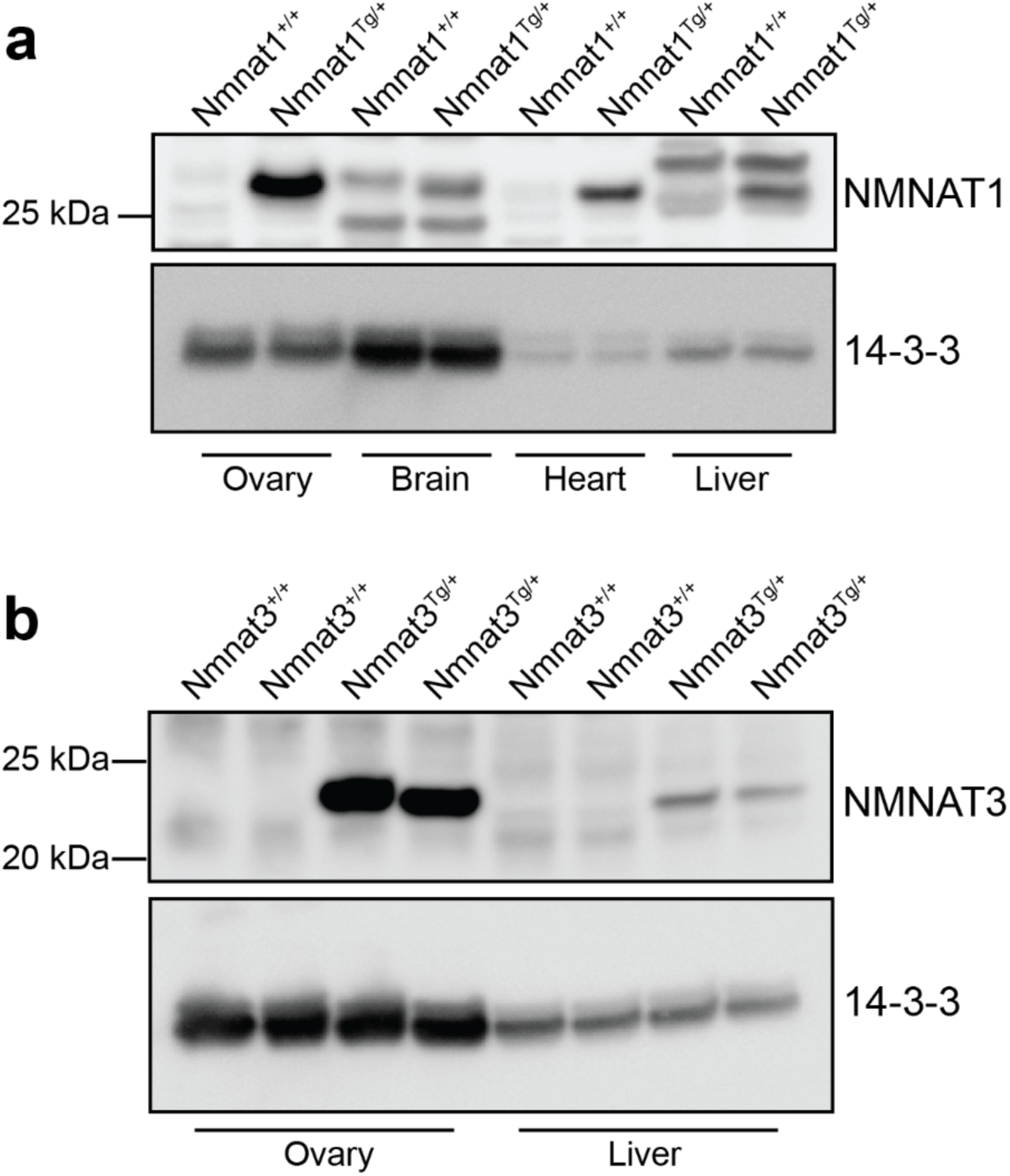
Transgenic overexpression of NMNAT1 and NMNAT3 in ovaries. Western blot for (**a**) NMNAT1 in the ovaries of *Nmnat1*^*Tg/+*^ animals, and (**b**) NMNAT3 in the ovaries of *Nmnat3*^*Tg/+*^ animals. Each lane represents tissue from a separate animal.

**Extended Data Figure 5.**
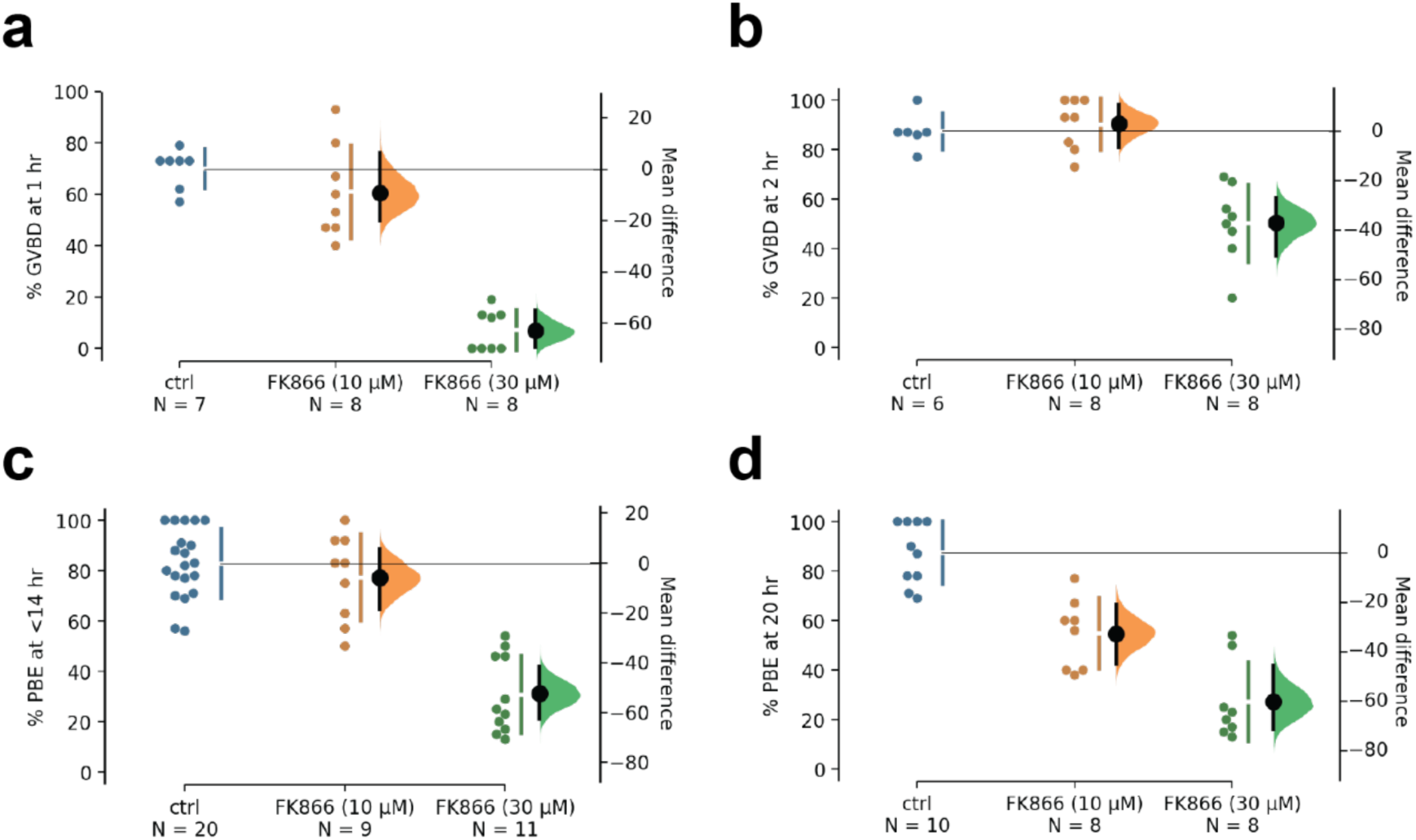
Inhibition of NAD synthesis slows meiotic progression. GV stage oocytes collected from 8-week old C57BL6 females were treated with the indicated concentrations of the NAMPT inhibitor FK866, and meiotic progression assessed. FK866 reduced germinal vesicle breakdown (GVBD) at (**a**) 1 hr and (**b**) 2 hr, as well as polar body extrusion (PBE) at (**c**) 14 hr and (**d**) 20 hr.

**Extended Data Figure 6.**
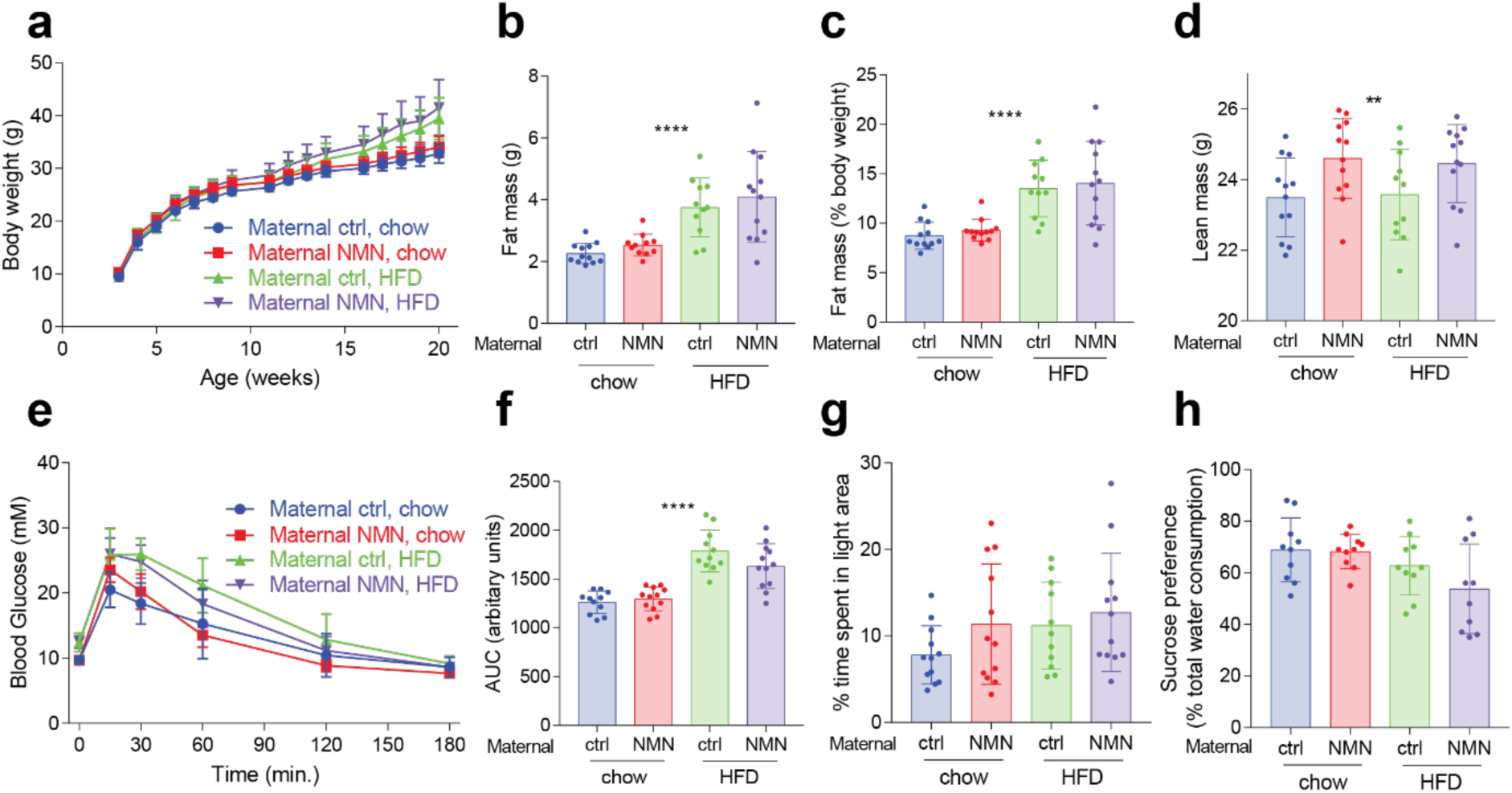
Maternal NMN treatment does not impact growth, metabolism or behaviour in offspring. Male offspring from females treated with or without NMN were maintained on standard chow or high fat diets (HFD). **a**, Body weight was measured on an ongoing basis. **b**, Fat mass and (**c**) lean body mass, also expressed as (**d**) body composition were assessed by quantitative MRI, with a difference in fat mass from HFD but no effect of maternal NMN treatment (\*\*\*\**p*<0.0001 diet effect, diet *F*=32), and a small but significant increase in lean body mass (\*\**p*=0.0054 maternal NMN effect, diet F=0.0096, maternal NMN F=8.604) with maternal NMN treatment. Metabolic homeostasis was measured by (**e**) glucose tolerance test, quantified by (**f**) area under the curve (AUC) (****p<0.0001, diet effect). Behaviour was assessed by the (**g**) light/dark box test for anxiety-like behaviour, and the (**h**) sucrose preference test for depression-like behaviour. 2-way ANOVA, *n*=11-12 per group. Data presented as mean ± s.d. Full statistical analyses in Extended Data Table 1.

**Extended Data Figure 7.**
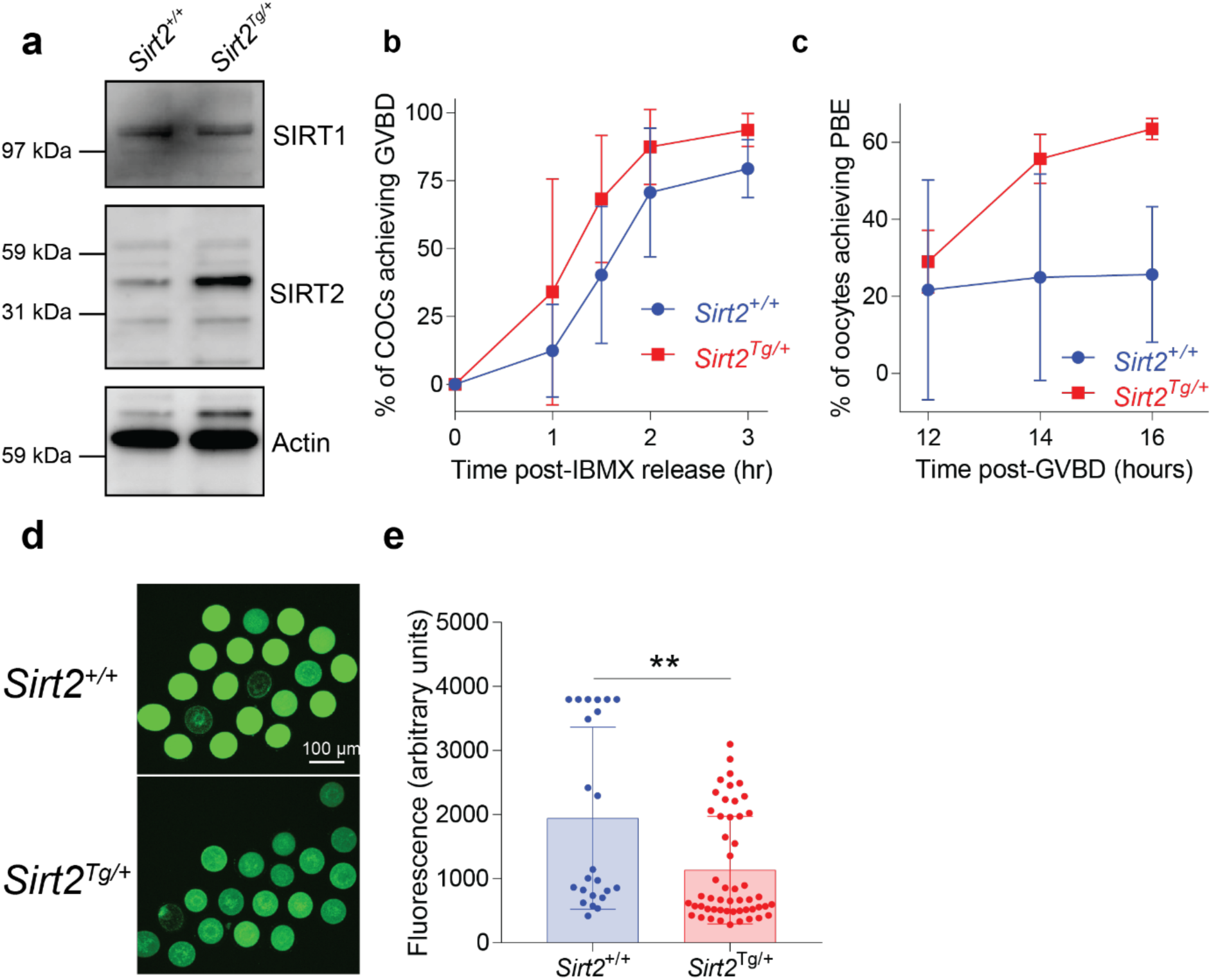
Data from Figure 4 presented using traditional null hypothesis statistics testing. **a**, Western blot for SIRT2 in oocytes from whole body *Sirt2* overexpressing transgenic mice (40 oocytes per lane). **b**, Germinal vesicle breakdown and (**c**) polar body extrusion rates in oocytes from 14-month old *Sirt2*^*Tg/+*^ or wild-type *Sirt2*^*+/+*^ littermates, at indicated timepoints (*n*=3-5 experiments). **d**, Oocytes from *Sirt2*^*Tg/+*^ animals have improved resistance to reactive oxygen species (ROS) challenge following H_2_O_2_ treatment compared to oocytes from wild-type *Sirt2*^*+/+*^ littermates, as measured by H_2_DCFDA staining, quantified in **e** (\*\**p*=0.0024, *n*=23-52 oocytes per group). Data are mean ± s.d. Further statistical detail in Extended Data Table 1.

**Extended Data Figure 8.**
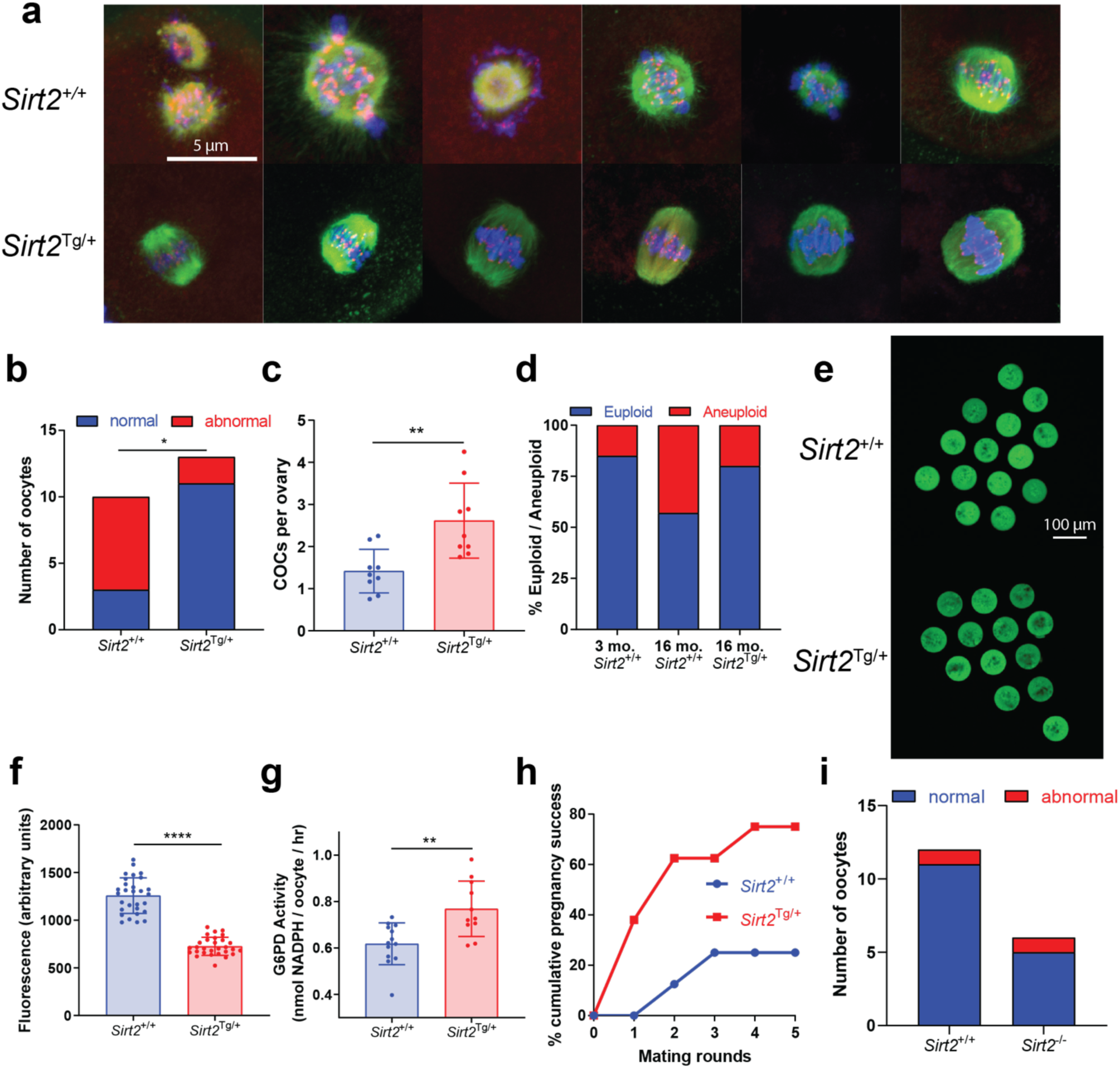
Data from main text Figure 4 presented using traditional null-hypothesis statistics testing. GV stage oocytes were recovered from 14 month-old *Sirt2*^Tg/+^ C57BL6 mice and matured in vitro to the MII stage, following which they were subjected to (**a**) Immunostaining for spindle assembly (β-tubulin in green, Hoescht for DNA in blue, kinetochores (ACA) in red), with (**b**) a quantitative improvement in spindle assembly in *Sirt2*^Tg/+^ transgenics compared to *Sirt*2^+/+^ littermates (n=10-13 oocytes per group). **c**, Oocyte yield from reproductively aged (14 month old) PMSG stimulated *Sirt2*^*Tg/+*^ and wild-type *Sirt2*^*+/+*^ littermates (\*\**p*=0.0030, *n*=9 animals per group). **d**, Aneuploidy rates in oocytes from young (2 month old) and aged (16 month old) *Sirt2*^*Tg/+*^ and wild-type *Sirt2*^*+/+*^ littermates (n=30 young wild-type, 7 *Sirt2*^+/+^ and 5 *Sirt2*^*Tg*/+^ oocytes from 4 animals per group). Oocytes from *Sirt2*^*Tg*/+^ mice had decreased ROS levels as determined by (**e**) H_2_DCFDA staining, quantified in (**f**) (\*\*\*\**p*<0.0001, *n*=29 oocytes from four 3-4 month old animals per group), due in part to (**g**) increased G6PD enzyme activity (**p=0.0019, n=11-13, each data point represents 5 pooled oocytes from 4 animals per group). **h**, Timed mating trials starting from 15 months of age to determine cumulative pregnancy rates, as determined by ultrasound imaging of a foetal heartbeat (χ^2^ test p=0.1319 after 5 mating rounds, n=8 animals per group). SIRT2 is sufficient, but necessary for oocyte integrity as (**i**) oocytes from *Sirt2*^*-/-*^ knockout animals display normal spindle assembly when as assessed as in (**a**). Data analysed by two-sided Fisher’s exact test in (**b**) and (**h**), two-tailed Student’s t-test in (**c**), (**f**) and (**g**). Data are mean ± 13 s.d; individual data points are shown. Full statistical analyses in Extended Data Table 1 and Supplementary Information.

**Extended Data Fig. 9.**
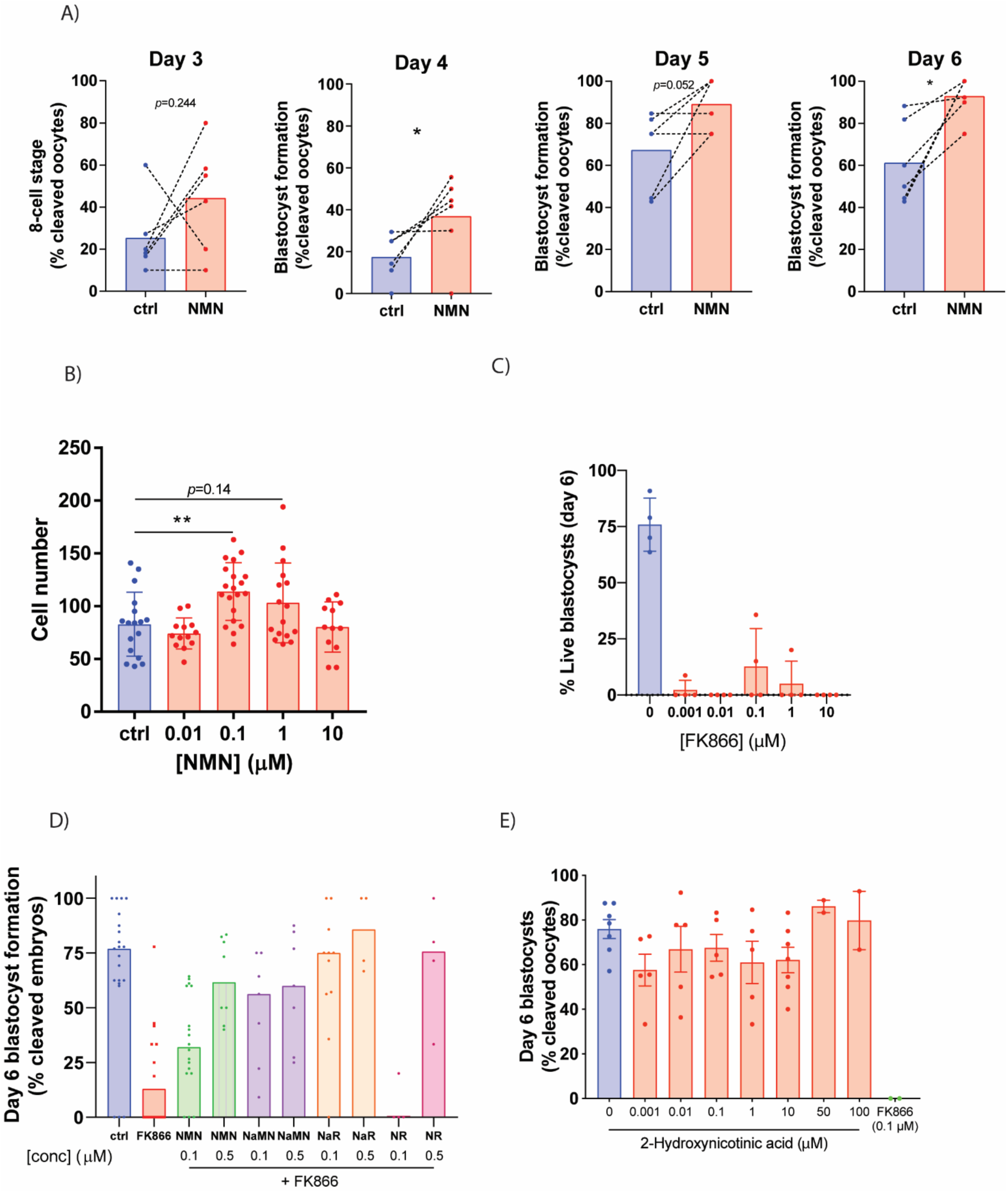
Data from Figure 5 presented using traditional null hypothesis statistics testing. MII oocytes from (**a**) aged (12 month old) mice were subjected to IVF, and post-fertilisation embryos maintained in media containing 1 μM NMN until day 6 of embryo development, with the percentage of embryos reaching developmental milestones (8-cell or blastocyst) shown for days 3-6. The addition of NMN could also enhance cell count at 92 hr in blastocysts from young animals matured in (**b**) simple defined media, which induces culture stress. **c**, Treatment with the NAMPT inhibitor FK866 at the indicated concentrations causes embryo death at day 6 (% surviving blastocysts shown), which (**d**) can be rescued by treatment with the NAD precursors NMN, nicotinic acid mononucleotide (NaMN), nicotinic acid riboside (NaR) and nicotinamide riboside (NR). **e**, Treatment with the NaPRT inhibitor 2-hydroxynicotinic acid has a minimal effect on blastocyst formation, compared to FK866 treatment (day 6 data shown). Data analysed in **a** by paired two-tailed t-test, in **b** by one-way ANOVA with Hold-Sidak multiple comparison test. Data are mean ± s.d. **p<0.01

**Extended Data Fig. 10.**
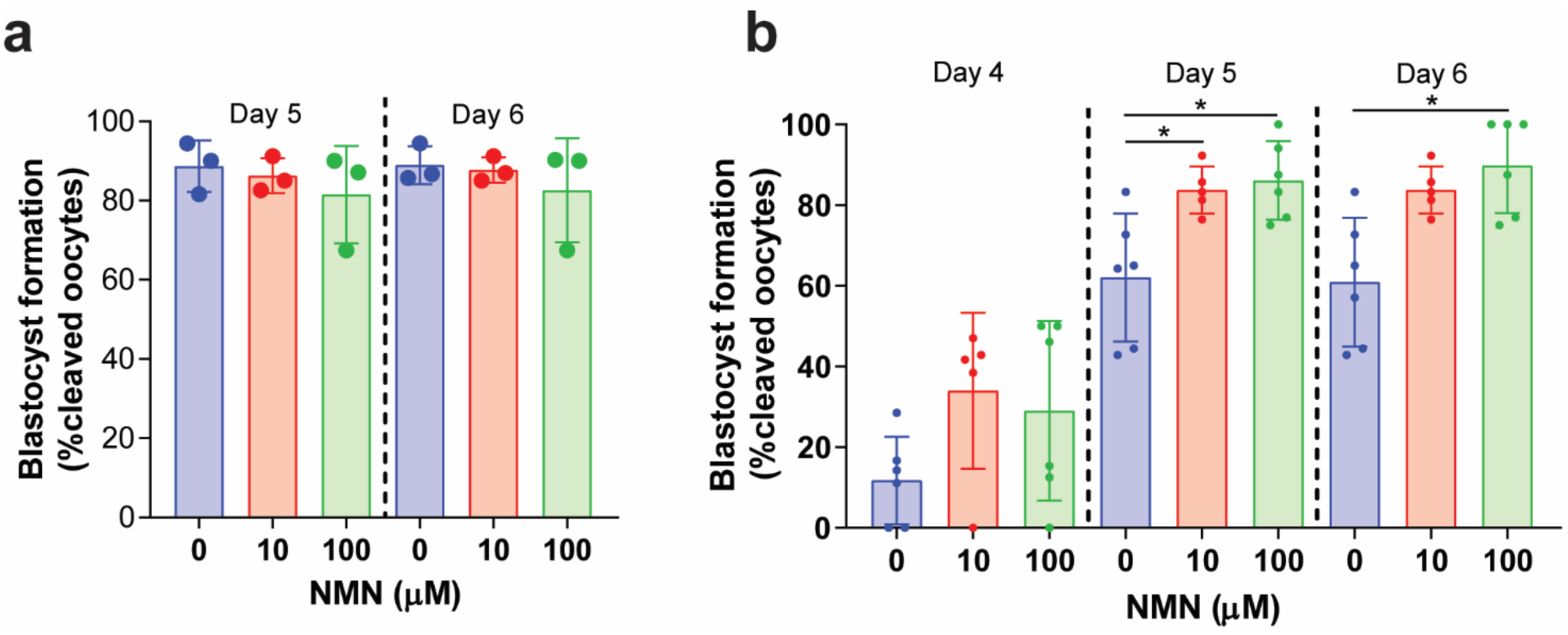
NMN enhances embryo formation from aged but not young females. Oocytes from (**a**) 4-week old or (**b**) 12-month old females were subjected to IVF, with the addition of NMN at indicated concentrations to embryo media. Blastocyst formation rates were assessed as indicated. Data analysed within each day by one-way ANOVA with Kruskal-Wallis multiple comparison. Day 5, 0 vs 10, *p*=0.0466; 0 vs 100, *p*=0.0132. Day 6, 0 vs 100, *p*=0.0100.

**Extended Data Fig. 11.**
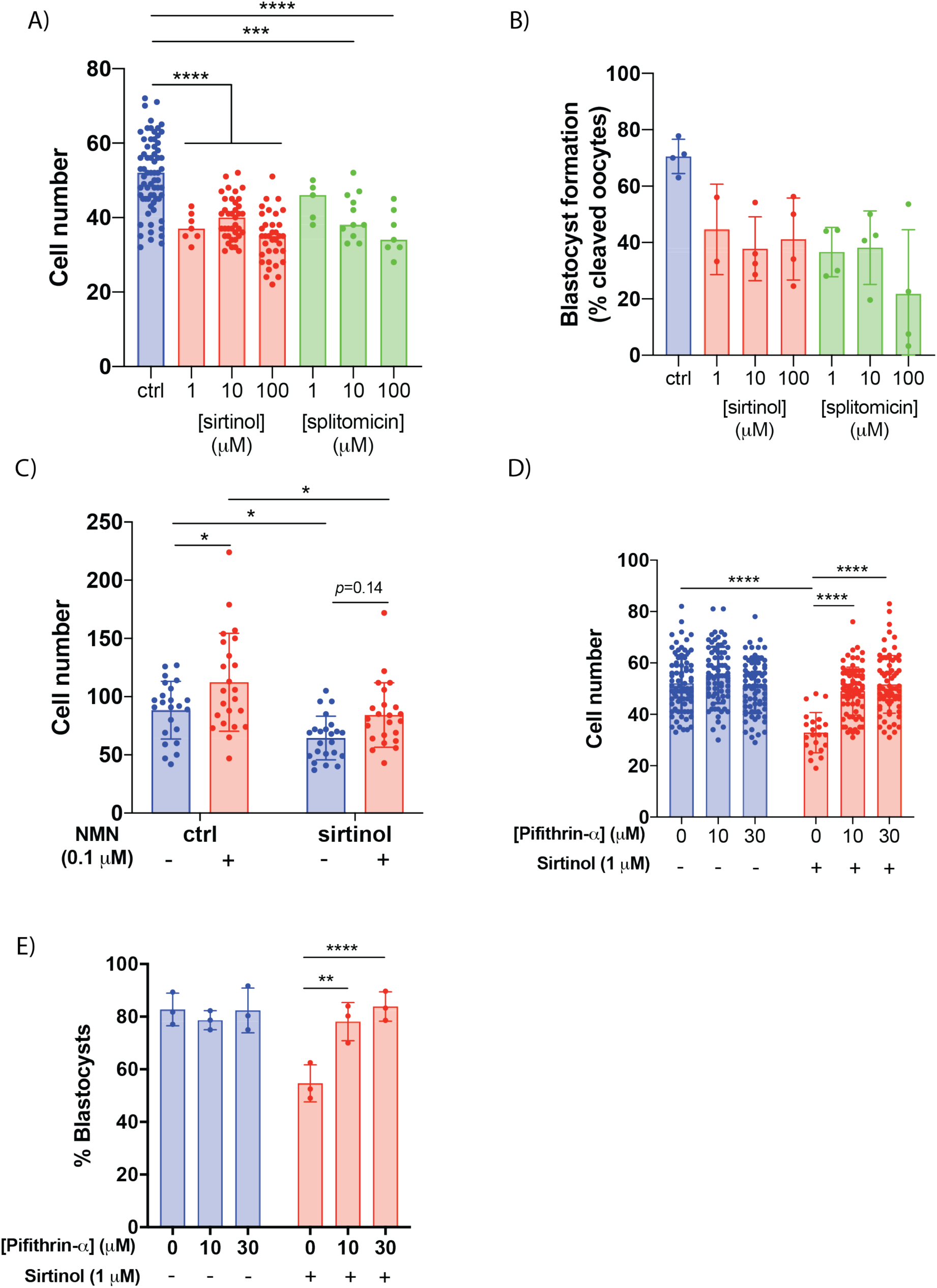
Data from main text Fig. 6 presented using traditional null hypothesis statistics testing. Treatment with the small molecule sirtuin inhibitors sirtinol or splitomicin (**a**) inhibits blastocyst formation, with (**b**) decreased cell count in those blastocysts that are formed. **c**, Co-treatment of sirtinol treated embryos with NMN rescues this reduction in cell count, indicating that the benefits of NMN are partially independent of sirtuins activity. Treatment with the p53 inhibitor pifithrin rescues (**d**) blastocyst formation and (**e**) cell count in embryos treated with sirtinol. Data from embryos maintained in simple defined media, fixed at 92 hr post-fertilisation. Data are analysed in **a, d, e** by one-way ANOVA with Holm-Sidak multiple comparison test, in **c** by 2-way ANOVA with Sidak’s multiple comparison test. *p<0.05, ****p<0.0001.

## Notes

† Denotes authors with involvement in Jumpstart Fertility Pty Ltd through direct employment, indirect salary support, sponsored research lab support, shareholding or directorship.

