## Supplementary Methods for "NAD^+^ repletion rescues female fertility during reproductive ageing"

Animals

Transgenic and wild-type strains were bred at Australian Bio-Resources (ABR) in Moss Vale, NSW Australia and delivered to UNSW Sydney at 6-7 weeks of age. Animals were maintained in individually ventilated cages at 22°C at 80% humidity at a density of up to 5 per cage, with *ad libitum* access to food and water. All water in this animal house was acidified to pH 3 with HCl to decrease microbial growth. The UNSW animal house maintained a 12 hr light/dark cycle with lights on at 0700 and off at 1900. Experiments were carried out with prior approval of the UNSW Animal Care and Ethics Committee (ACEC) under ACEC numbers 13/134B, 15/134A and 18/133A. UNSW ACEC operates under animal ethics guidelines from the National Health and Medical Research Council (NHMRC) of Australia. The generation of SIRT2, NMNAT1 and NMNAT3 transgenic mouse strains used here were as previously described(1, 2).

Ovarian stimulation, oocyte collection for *in vitro* maturation and spindle assessment (Fig. 1, 2)

To collect germinal vesicle stage cumulus oocyte complexes (COCs) for counting and for in vivo maturation for spindle morphology assessment, both ovaries from each mouse were recovered 44 to 46 hours after ovarian stimulation with 7.5 IU Pregnant mare's serum gonadotropin (PMSG) (MSD Animal Health, Australia), which was administered by intraperitoneal (i.p.) injection. Fat and tissue surrounding the ovaries was carefully and rapidly removed under the microscope. Oocytes were mechanically released from ovaries under M2 medium supplemented with 3 mg/ml bovine serum albumin (BSA) in the presence of the growth inhibitor 3-isobutyl-1-methylxanthine (IBMX) (Sigma Aldrich, Australia) at 50 μM using 27-guage needle. The released oocytes, including cumulus oocyte complexes (COC) and denuded oocytes (DO) in the germinal vehicle (GV) stage were collected by aspirator tubes (Sigma Aldrich) and transferred into another drop of IBMX-M2 media overlayed with paraffin oil (EMD Millipore corporation, USA), which was pre-warmed to 37°C. Oocytes were then counted, and the cumulus cells removed by mechanical disruption with a mouth pipette. Oocytes were then rinsed 3 times in IBMX-free M2 (Sigma Aldrich, Australia) containing 3 mg/ml BSA, following 24 hours pH calibration in 5% $CO_2$, to clear off the IBMX from medium, then transferred into another clean, pH calibrated M16 medium that was overlayed with oil and returned to the incubator under 37°C in a humidified atmosphere of 5% $CO_2$ to assess subsequent meiotic development. Meiotic maturation measurements included the rates of GVBD and first polar body extrusion (PBE). The rates of GVBD were measured by counting the number of oocytes where there was a disappearance of nuclear membranes in the first two hours after release. Oocytes that failed to develop into the GVBD stage after 2 hours were discarded, and the remaining GVBD stage oocytes were then continuously incubated to assess PBE development, indicated by the production of first polar body, in 14 hours, 16 hours and 20 hours afterward.

For immunofluorescence and spindle analysis, oocytes which had achieved polar body extrusion were removed from culture medium and very briefly washed through PHEM solution (60 mM PIPES at pH 6.9, 25 mM HEPES, 10 mM EGTA, 2 mM $MgCl2.7H_2O$), prior to being pre-permeabilized in 0.25% Triton X-100 (Sigma) in PHEM for 5 seconds at room temperature to soften the zona pellucida. Oocytes were then fixed in 3.7% paraformaldehyde (Sigma) in PHEM for 20 minutes before being permeabilized for 10 minutes in 0.25% Triton X-100 in PBS. Oocytes were then washed in PBS containing 0.5% Bovine serum albumin (BSA) (Sigma) (wash solution) for 5 minutes at room temperature before being blocked overnight in PBS containing 3% BSA and 0.05% Tween-20 (blocking solution) at 4oC. The following morning, oocytes were allowed to equilibrate to room temperature and were probed with primary antibodies.

For immunolabelling, human anti-centromere antibodies (ACA; 1:40; ImmunoVision) and; mouse anti-β-tubulin (1:800; Sigma) were used to label kinetochores and microtubules, respectively. Following primary antibody incubation, the oocytes were then washed and placed into the following secondary antibodies: Alexa Fluor 546-labelled goat anti-human (1:200; Invitrogen) for detecting ACA; and Alexa Fluor 488-labelled goat anti-mouse (1:200; Invitrogen) for detecting β-tubulin. DNA was labelled using 60-second incubation in Hoechst 33342 (1 μg/ml; bis-benzimide; Sigma) in PBS at RT. Oocytes were then transferred to 1-2 μl micro-drops of PBS under mineral oil in glass bottom dishes for confocal imaging.

Serial Z sections of fixed oocytes were captured at 0.5 μm intervals, to span the entire region of the spindle, using an A1 MP+ multiphoton confocal microscope (Nikon, Japan) equipped with a C-Apochromat 63x/1.4 NA oil immersion objective, and processed using NIS-Elements C acquisition and analysis software (Nikon, Japan). Confocal image stacks were projected and further processed using the imaging software FIJI.

For spindle analyses, oocytes exhibiting barrel-shaped bipolar spindles with well-organized microtubule fibers, along with tightly aligned chromosomes on the metaphase plate, were considered as normal.

Monastrol treatment and ploidy assessment

Following in-vitro maturation, MII oocytes were cultured for 2 h in M2 medium (Sigma) containing 100 μM monastrol (Sigma), a kinesin-5 inhibitor used to align chromosomes for investigations of chromosome number and dynamics (3). Oocytes were then fixed, blocked and immunostained as described in the above section. Kinetochores were detected using human anti-centromere primary antibody (ACA; ImmunoVision), followed by incubation in Alexa Fluor 546-labelled goat anti-human (1:200; Invitrogen). DNA was labelled using 10-minute incubation in a 1:5000 dilution of Sytox Green nucleic acid stain (Invitrogen). Images were acquired at 0.5 μm intervals using the microscope described above and processed using the FIJI software.

To obtain a chromosome count in each oocyte, the Z-projection image was initially analyzed to determine the number of kinetochore pairs. Where there was possible overlap of kinetochores or equivocation, serial confocal sections were analyzed while carefully noting the orientation of the chromosomes. Oocytes with excessive chromosome clumping and those with ambiguous counts were excluded from the final aneuploidy assessment. A single trained observer performed all calculations and ploidy determination.

Oocyte collection and confocal imaging – DCFDA staining

To collect mature MII COCs, mice were super-ovulated with an intraperitoneal (i.p.) injection of 10 IU pregnant mare serum gonadotrophin (Folligon; Intervet, Boxmeer, Holland) to stimulate follicle growth, followed by a 10 IU i.p. injection of human chorionic gonadotrophin (hCG; Chorulon, MSD Animal Health, Australia) 46 hr later to induce ovulation. COCs were collected form oviductal ampullae using a 27-guage needle and collected in HEPES-buffered α-minimum essential medium (α-MEM; Gibco Life Technologies, Grand Island, NY) supplemented with 3 mg/ml bovine serum albumin (BSA; Sigma Aldrich, St Louis, MO) 14 – 16 h after hCG injection.

In order to assess the response of the oxidative stress defence mechanisms of oocytes, intra-oocyte glutathione (GSH) and reactive oxygen species (ROS) levels were assessed using the fluorescent probes 5-(and-6)-Chloromethyl-2',7'-dichlorodihydrofluorescein diacetate (CM-$H_2$DCFDA) (Invitrogen). In unchallenged oocytes as shown in Fig. 1e, as previously described (4), MII oocytes were mechanically stripped of their cumulus cells prior to staining and imaging. Oocytes were incubated in HEPES-buffered buffered α-MEM containing 10 μM CM-$H_2$DCFDA for 30 mins at 37°C protected from light. Oocytes were washed three times in HEPES-buffered α-MEM medium before transfer into a 2 μL drop of HEPES-buffered α-MEM medium, overlayed with paraffin oil in a glass bottom 35 mm confocal dish (MatTek Corp.). Oocytes were imaged immediately under bright field and fluorescence using a Nikon A1+ confocal microscope using a 20X objective at 37°C. For $H_2O_2$ challenged oocytes as shown in Extended Data Fig. 1d, MI cumulus oocyte complexes were recovered from PMSG stimulated animals and mechanically denuded, and incubated overnight in α-MEM with 50 μM IBMX in a 5% $CO_2$ incubator at 37°C. Oocytes were then incubated in HEPES buffered α-MEM with IBMX with the addition of 25 μM H2O2, then washed three times and allowed to recover in HEPES buffered α-MEM with IBMX for 90 min, and stained for 30 min in HEPES buffered α-MEM with IBMX containing 10 μM DCFDA. Oocytes were then washed and imaged immediately by confocal microscopy at 37°C. ROS-induced cleavage of CM-$H_2$DCFDA to DCF fluorescence was examined using an excitation and emission wavelength of 488 nm and 520 nm. Images were taken under fixed conditions with respect to laser energy, signal detection (gain) and pinhole size. Confocal z-stacks of 10 μm thickness with 1 μm increments between slices were recorded at the equatorial plain of the oocytes. Mean relative fluorescence intensities of DCF were measured in a minimum of 25 oocytes per group using ImageJ Fiji software (5).

IVF and embryo culture

Oocyte developmental competence was evaluated by examining the capacity of oocytes to support preimplantation embryo development following *in vitro* fertilization (IVF). IVF and embryo culture were performed as previously described by Yeo et al (6). All IVF media was purchased from IVF Vet Solutions (Adelaide, Australia). Briefly, following stimulation of females with 5 IU (young) or 10 IU (aged) PMSG and hCG, COCs were collected form oviductal ampullae using a 27-guage needle and collected in HEPES-buffered α-minimum essential medium COCs were washed three times in wash media (Research Wash) supplemented with 4mg/ml of BSA and co-incubated with capacitated sperm in fertilisation media supplemented with 4mg/ml of BSA for 4h at 37°C with 5% $O_2$, 6% $CO_2$ and 89% $N_2$. Sperm was recovered from hybrid CBB6F1 males > 12 weeks of age. Presumptive zygotes were washed three times in wash media and then once in cleavage media (Research Cleave) supplemented with 4 mg/ml BSA and cultured in drops over-layered with mineral oil at a density of 1 embryo per 2 µl. Fertilisation rates and subsequent on-time development were assessed every 24 hours.

In the case of simple defined media growth conditions (Fig. 5b, Fig. 6), oocytes were collected as above, and then maintained in human tubal fluid supplemented with glutamine and EDTA (HTF-GE) media further was supplemented with 3 mg/ml BSA, which is a minimal essential media that does not contain any hormones or trophic ligands (7). Embryos were cultured individually in 10 µL drops to further induce culture stress from diluting autocrine trophic ligands. Embryos were fixed at 92 hr post-fertilisation for cell counting.

<u>G6PD activity</u>

MI oocytes from PMSG stimulated animals were recovered from ovaries and mechanically denuded of the cumulus layer via mouth pipetting, and collected into 50 mM Tris-HCl pH 7.44 buffer in a volume of 35 µL with 5 oocytes per sample. The oocytes were lysed following a single freeze-thaw cycle, and spun down at 16,000g, 4 degrees for 10 minutes to separate cell debris. The supernatant (34 µL) was subjected to measurement for glucose-6-phosphate dehydrogenase (G6PD) activity, which was determined as described previously (8). Both 6-phosphogluconate dehydrogenase (6PGD) activity and G6PD+6PGD activity were measured separately, and endogenous G6PD activity was determined by subtracting the 6PGD activity alone from the G6PD+6PGD total enzyme activity value. Each reaction mixture consists of 50 mM Tris-HCl, pH 7.44, 2 mM $NADP^+$ and sample in which G6PD activity was to be measured. Reactions were initiated with the addition of 4 mM G6P + 4 mM 6PGL or 4 mM 6PGL alone and activity was measured using a spectrophotometer at room temperature, in a total volume of 60 µL in a 384 well plate. NADPH production was measured at absorbance 340nm every 30 seconds, and the rate of NADPH synthesis determined as a unit of AU/hour.

### NAD$^+$ assay

NAD$^{\pm}$ levels as shown in Fig. 1 were assayed based on the previously described NAD$^{\pm}$ cycling method of Zhu and Rand (9). Following euthanasia, ovaries were rapidly dissected out of animals and snap frozen in liquid nitrogen, then stored at -80°C. Samples were then homogenised in NAD$^{\pm}$ extraction buffer (10 mM nicotinamide, 50 mM Tris HCl, 0.1% Triton X-100, pH 7.4) by sonication. Samples were then centrifuged at 7,000 g for 5 min at 4$^o$C to remove insoluble debris. Aliquots were taken for later protein assay, and samples were then passed through 10 kDa Amicon filters at 14,000 g, 30 min at 4$^o$C to remove proteins from the sample. Each sample was measured in technical triplicate, with 25 µL sample added to 100 µL ADH cycling mix (0.2 mg/ml alcohol dehydrogenase enzyme, 2% ethanol, 100 mM Tris pH 8.5). Samples were allowed to cycle for 10 min at room temperature, followed by 50 µL addition of an MTT/PMS solution (0.1 mM phenazine methosulfate, 0.8 mM 3-(4,5-Dimethylthiazol-2-yl)-2,5-diphenyltetrazolium bromide), 100 mM Tris-HCl pH 8.5). Plates were then incubated for 15 min and absorbance was measured at 570 nM NAD concentrations were extrapolated from a standard curve and normalised to protein concentrations determined by BCA protein assay.

### Hyperspectral imaging

In this work we utilised a fluorescence microscope adapted for multispectral imaging by adding a number of custom-made fluorescent channels (10-13) to characterise native fluorescence of oocytes. Images were taken in the following channels as described in Supplementary Table 1. We generated a sequence of autofluorescence images of the same cells in each of these channels at 40 times magnification (40x). To acquire the fluorescence images, a sensitive camera was used (Prime 95B cMOS from Photometrics). The camera was operated below -65°C to reduce sensor-induced noise.

To prepare spectral images for quantitative analysis and spectral unmixing, firstly, a multistep image preparation procedure was carried out (10, 11, 14, 15). This procedure removes sources of errors, reduces noise and standardizes the spectral images. These steps include (i) image equalization, (ii) denoising, (iii) flattening of background illumination, and (iv) cell segmentation which was reported in detail in our previous studies (15). Briefly, to equalize the images, the intensity value at each channel is converted into the units of photons per pixel per second (PPS). PPS unite standardizes the images acquired with different acquisition parameters such as camera quantum efficiency and exposure time. Next, to reduce random noises such as spikes and Poissonian noise, a 'threshold limiting window' and wavelet filter were employed, respectively.

Further, two reference multispectral images including water and calibration were captured using our multispectral microscope system at the beginning of each experiment. After denoising, the water image was subtracted from the spectral images to remove the unavoidable autofluorescence signals from the microscope slide, petri dishes, dirt on sensors etc which forms an additive autofluorescence background contribution to all spectral images. The microscope was calibrated using a fluorimeter (FluoroMax Plus-C, Horiba). To this aim we used the calibration fluid which is a mixture of 30 μM NADH (quantum yield 0.019) and 5μM riboflavin (quantum yield 0.24) with non-zero fluorescent response across all our spectral channels. The fluorescence spectrum of this fluid is measured using a fluorimeter and imaged on the multispectral microscope in all our spectral channels. The fluorescence images of the calibration fluid were then smoothed and associated random noises were minimised. The smoothed spectral images with subtracted water background, were then divided by the calibration fluid image and multiplied by fluorescence response of the calibration fluid measured by fluorometer. This was done on a channel by channel basis. Finally, to perform the unmixing process at a single cell resolution, the cells were isolated manually (segmented) by using their DIC image superimposed on selected autofluorescence images.

In the unmixing process the measured spectral characteristics of the cells were compared to the reference spectra of specific fluorophores to find their abundance. In this study, unmixing was performed sing a linear mixing model (LMM). The LMM describes the fluorescent signal of each pixel is a linear combination of a fraction of endmember component spectra. LMM defines weights associated with the concentration of the molecules corresponding to these component spectra. These concentrations were presented as abundance fractions (16-18) and the Robust Dependent Component Analysis (15) (RoDECA) algorithm was employed to identify the dominant native fluorophores and calculate their relative abundance.

| Spectral channel | Excitation wavelength (nm) | Emission wavelength (bandwidth) (nm) | Dichroic mirror long pass (nm) | Power at objective(µW) | Exposure time (s) |
|---|---|---|---|---|---|
| 1 | 345±5 | 414 (46) | 389 | 6.0 | 10 |
| 2 | 345±5 | 451 (106) | 389 | 6.2 | 6 |
| 3 | 345±5 | 575 (59) | 552 | 5.7 | 10 |
| 4 | 358±5 | 414 (46) | 389 | 5.2 | 10 |
| 5 | 371±5 | 414 (46) | 389 | 6.8 | 10 |
| 6 | 358±5 | 451 (106) | 389 | 5.2 | 10 |
| 7 | 371±5 | 451 (106) | 389 | 6.9 | 10 |
| 8 | 358±5 | 575 (59) | 552 | 6.3 | 10 |
| 9 | 371±5 | 575 (59) | 552 | 9.9 | 10 |
| 10 | 377±5 | 575 (59) | 552 | 18.6 | 10 |
| 11 | 437±5 | 575 (59) | 552 | 28.8 | 10 |
| 12 | 457±5 | 575 (59) | 552 | 25.2 | 10 |
| 13 | 476±5 | 575 (59) | 552 | 24.1 | 10 |
| 14 | 358±5 | 594 (long-pass) | 552 | 6.7 | 10 |
| 15 | 371±5 | 594 (long-pass) | 552 | 10.4 | 10 |

**Supplementary Table 1. spectral specification of channels used in this study.**

High fat feeding

Animals were maintained on chow or high fat diets as described previously (19). Chow diet (Gordon's Specialty Stock Feeds, Yanderra, New South Wales, Australia) comprised 8% calories from fat, 21% calories from protein, and 71% calories from carbohydrate, with total energy density of 2.6 kcal/g. High-fat diet (HFD) was prepared in-house and comprised 45% calories from fat (beef lard), 20% calories from protein, and 35% calories from carbohydrate at a density of 4.7 kcal/g, based on rodent diet D12451 (Research Diets, New Brunswick, NJ, USA).

Offspring studies

Male offspring from the control and NMN only treated groups of the breeding trials shown in Fig. 3A were maintained to determine whether maternal NMN exposure adversely impacted offspring health. NMN exposure in the mothers was ongoing in drinking water (2 g/L) from 16 weeks of age, including during mating, pregnancy and lactation. Offspring were weaned onto normal drinking water without NMN, and either standard chow diet or high fat diet. Animals were subjected to body composition analysis and glucose tolerance testing at 10 weeks of age, and behavioural analyses at 5 months of age.

Sucrose Preference Test (SPT)

Sucrose Preference Test is a standard test for assessing anhedonic behavior, a symptom of depression. This test was performed at 5 months of age in male offspring of NMN treated females according to a previous study with some modifications (20). Mice were housed individually in separate cages with access to two bottles. Mice were firstly pre-exposed to sucrose solution (1%, w/v) for 24 h as a training to adapt to the newly introduced sucrose solution. They were given access to 2 bottles (sucrose 1%, tap water). On the second day, food and water were removed for 6 hours before being giving access again to sucrose and water. In order to avoid place preference, the position of the bottles was changed, and mice were given to both 1% sucrose solution and water for 24 h. Sucrose preference was calculated according to the following formula: sucrose preference (%) = sucrose intake (g) / [sucrose intake (g) + water (g)] x 100%

Light/Dark Test

The light/dark test (LDT) is a standard test to assess anxiety-like behaviour. This test was conducted during light phase at 5 months of age according to a previous protocol (20). The LDT apparatus comprised of bright and dark perspex compartments (24 cm 24 cm 27 cm) connected by a small opening (10 cm 10 cm). The light reading on the floor of the light area was 4000—5000 lux and the dark area was lit between 1 and 10 lux. Mice were gently placed in the light area, facing away from the doorway. The total time spent in light and dark areas were recorded for 5 min. The percentage of time spent in light was calculated using the following formula: % time in light area = time spent in light /[(time spent in dark + time spent in light)].

Glucose tolerance test (GTT)

GTT was as described previously (19). Animals were fasted from 0800 on the day of the test. At 1400, baseline blood glucose was assayed using a hand-held glucose testing meter (Accu-Check, Roche) through a scalpel nick less than 1 mm from the tip of the tail, with tails protruding from a small upturned cardboard box used to calm animals. Animals were injected with 25% glucose (i.p. injection) at 2 g/L lean body mass, and glucose was assessed at indicated timepoints from tail bloods. Lean body mass was assessed by quantitative MRI.

Body composition measurements

Body composition was determined by quantitative MRI using an echoMRI (Houston, TX USA) in conscious animals.

Oral gavage and pharmacodynamics

Animals were administered a single oral gavage of NMN in a volume of 100 uL water per 20 g body mass for a total dose of 500 mg/kg. Animals were euthanased at the indicated timepoint, and ovaries rapidly dissected, cleared of surrounding adipose tissue, and snap frozen for subsequent analysis of $NAD^+$ levels.

Breeding trial design

The reproductive capacity of 16-month old SIRT2-Tg mice (n=8) were compared to that of age-matched wild-type littermates (n=8). We employed a timed-mating protocol, whereby each breeding round commenced with group-housed female mice being ‘scented’ using dirty bedding from a male’s cage for 3 days. This design allowed for stimulation and synchronization of the females’ estrous cycles through the Whitten Effect (21). Each female was then placed into an individually-housed proven male breeder’s cage in a 1:1 ratio (‘co-habitation’). This transfer was made in the evening, in anticipation of night-time ovulation (2). The animal-holding facility default light setting is a 12-hour light/12-hour dark cycle. The following early morning each female would be assessed for the presence of viscous vaginal plug as evidence of copulation and, regardless of plug status, were separated to prevent daytime mating. Females with no plug would be replaced with the same male the following evening to undergo another night of mating. If, following 3 consecutive nights of co-habitation, no plug had been observed, females would be separated and observed for weight gain for 5 days to account for possibility of missed plug, before undergoing re-scenting. A female would repeat the current mating round until a positive plug was confirmed. Females with positive plug were considered to have passed that mating round and would be housed separately before undergoing micro-ultrasound (VisualSonics Vevo 2100 Ultrasound) on day 15 post-copulation (pc) to determine pregnancy state, as determined by the presence of a foetal heartbeat. Females that had copulated, as demonstrated by plug presence, but did not fall pregnant would be deemed to have failed that mating round and would commence the next mating round with a different male stud, and so on. The data presented in Fig. 1h show the cumulative rate of pregnancy following confirmation of mating as described above.

Taking into account the high prevalence of labour difficulty and maternal cannibalism that was encountered in aged nulliparous mice, we were unable to determine offspring number from each pregnancy in the aged *Sirt2* transgenic cohort.

The breeding trial for NMN treated animals as presented in Fig. 2j was as above with some slight modifications. Younger animals were able to deliver litters without the above issues of labour difficulties or maternal cannibalism, and offspring were maintained with females until weaning at 21 days, after which females were returned to additional mating rounds, for a total of 6 mating rounds per animal. Animals were treated with NMN (drinking water, 2 g/L) from the age of 10 weeks, and then subjected to timed breeding at the age of 18 weeks, for 5 rounds with 7-8 weeks between rounds until the age of 50 weeks. The number of pups per litter were recorded and presented in Fig. 2j.

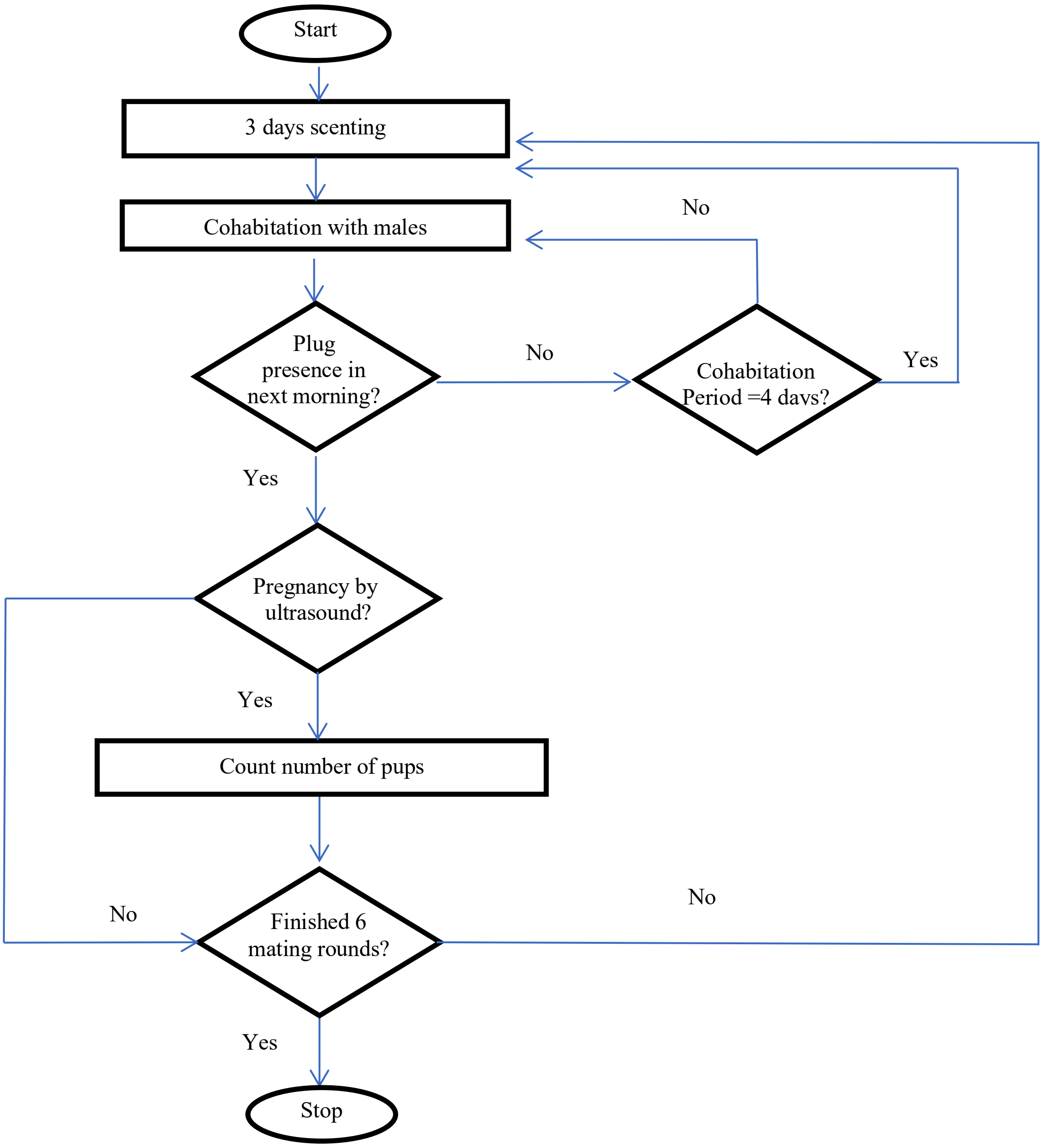


**Methods Figure 1.:** Experimental design of mating trials. Females were scented with soiled bedding from males for 3 days before cohabitation with males of proven breeding ability. Female mice were then separated from males if a vaginal plug was observed within 4 days of cohabitation. Otherwise, the maximum cohabitation period of 4 days was allowed, followed by separation regardless of the presence of a plug. Foetal heartbeat, as determined by micro-ultrasound imaging, was then used as an indicator of pregnancy after 21 days of cohabitation by ultrasound. Non-pregnant mice were then returned to the breeding process. Mating rounds were terminated after 6 rounds were completed, or unless animals had to be censored from the study due to severe illness.

### Statistics

Prior to conducting studies, power calculations were performed to determine sample size using G*Power 3.1, based on effect sizes observed in preliminary pilot studies.

There are strong arguments to move beyond null-hypothesis statistics tests (NHST) that make dichotomous decisions of significance thresholds based on a *p*-value (22). An alternative that has been proposed is the use of estimation statistics (23), which provides several advantages. Based on these arguments, we have analysed our data using estimation statistics (24), which were used to present figures in the main text. We recognise however that this is not yet a widely used form of statistics, and to allow interpretation from readers who are not familiar with this system, we have also provided traditional NHST based analysis for each Figure of this manuscript in Extended Data.

#### *Estimation graphs*

Data were analysed and figures produced using the DABEST (data analysis with boot-strap-coupled estimation) package for estimation statistics (24).

#### *Null hypothesis statistics testing (NHST) analysis*

For all variable outcomes, data sets were subjected to the D'Augusto & Pearson normality omnibus test. If all groups passed this test, data were analysed as either a two-tailed Student's t-test (2 groups), ANOVA (3 or more groups) or as a two-way ANOVA with a 2x2 study design, between group comparisons made by Holm-Sidak's multiple comparison test, as indicated in the text of figure legends. Non-parametric data sets were analysed by a two-tailed Mann-Whitney U-test (2 groups) or a Kruskal-Wallis test (3 or more groups) with between group comparisons made by Dunn's multiple comparison test. For data sets with binary outcomes, a two-sided Fisher's exact test was used. For tumour growth (Fig. 4b) data were analysed by both the Log-rank (Mantel-Cox) test for survival of time from start of therapy to euthanasia.

All data are presented as mean values with error bars representing standard deviation.

Statistical analyses were performed using GraphPad Prism version 7.04.

Detailed statistical calculations are available as .xml files in accompanying Supplementary Information.

### Supplementary methods references

1. North BJ*, et al.* (2014) SIRT2 induces the checkpoint kinase BubR1 to increase lifespan. *EMBO J* 33(13):1438-1453.
2. Yahata N, Yuasa S, & Araki T (2009) Nicotinamide mononucleotide adenylyltransferase expression in mitochondrial matrix delays Wallerian degeneration. *J Neurosci* 29(19):6276-6284.

3. Kapoor TM, Mayer TU, Coughlin ML, & Mitchison TJ (2000) Probing spindle assembly mechanisms with monastrol, a small molecule inhibitor of the mitotic kinesin, Eg5. *J Cell Biol* 150(5):975-988.
4. Li HJ*, et al.* (2016) Extending prematuration with cAMP modulators enhances the cumulus contribution to oocyte antioxidant defence and oocyte quality via gap junctions. *Hum Reprod* 31(4):810-821.
5. Schindelin J*, et al.* (2012) Fiji: an open-source platform for biological-image analysis. *Nat Methods* 9(7):676-682.
6. Yeo CX, Gilchrist RB, Thompson JG, & Lane M (2008) Exogenous growth differentiation factor 9 in oocyte maturation media enhances subsequent embryo development and fetal viability in mice. *Hum Reprod* 23(1):67-73.
7. O'Neill C (1997) Evidence for the requirement of autocrine growth factors for development of mouse preimplantation embryos in vitro. *Biol Reprod* 56(1):229-237.
8. Tian WN*, et al.* (1998) Importance of glucose-6-phosphate dehydrogenase activity for cell growth. *J Biol Chem* 273(17):10609-10617.
9. Zhu CT & Rand DM (2012) A hydrazine coupled cycling assay validates the decrease in redox ratio under starvation in Drosophila. *PLoS One* 7(10):e47584.
10. Gosnell ME*, et al.* (2016) Quantitative non-invasive cell characterisation and discrimination based on multispectral autofluorescence features. *Scientific reports* 6:23453 %@ 22045-22322.
11. Gosnell ME, Anwer AG, Cassano JC, Sue CM, & Goldys EM (2016) Functional hyperspectral imaging captures subtle details of cell metabolism in olfactory neurosphere cells, disease-specific models of neurodegenerative disorders. *Biochimica et Biophysica Acta (BBA)-Molecular Cell Research* 1863(1):56-63 %@ 0167-4889.
12. Rehman AU*, et al.* (2017) Fluorescence quenching of free and bound NADH in HeLa cells determined by hyperspectral imaging and unmixing of cell autofluorescence. *Biomedical optics express* 8(3):1488-1498 %@ 2156-7085.
13. Habibalahi A, Bala C, Allende A, Anwer AG, & Goldys EMJTOS (2019) Novel automated non invasive detection of ocular surface squamous neoplasia using multispectral autofluorescence imaging.
14. Vidal M & Amigo JM (2012) Pre-processing of hyperspectral images. Essential steps before image analysis. *Chemometrics and Intelligent Laboratory Systems* 117:138-148 %@ 0169-7439.
15. Mahbub SB, Plöschner M, Gosnell ME, Anwer AG, & Goldys EM (2017) Statistically strong label-free quantitative identification of native fluorophores in a biological sample. *Scientific reports* 7(1):15792 %@ 12045-12322.
16. Keshava N & Mustard JF (2002) Spectral unmixing. *Signal Processing Magazine, IEEE* 19(1):44-57.
17. Keshava N, Kerekes JP, Manolakis DG, & Shaw GA (2000) Algorithm taxonomy for hyperspectral unmixing. pp 42-63.
18. Keshava N (2003) A survey of spectral unmixing algorithms. *Lincoln Laboratory Journal* 14(1):55-78.
19. Wu LE*, et al.* (2014) Systemic VEGF-A neutralization ameliorates diet-induced metabolic dysfunction. *Diabetes* 63(8):2656-2667.
20. Maniam J & Morris MJ (2010) Palatable cafeteria diet ameliorates anxiety and depression-like symptoms following an adverse early environment. *Psychoneuroendocrinology* 35(5):717-728.
21. Jemiolo B, Harvey S, & Novotny M (1986) Promotion of the Whitten effect in female mice by synthetic analogs of male urinary constituents. *Proc Natl Acad Sci U S A* 83(12):4576-4579.
22. Amrhein V, Greenland S, & McShane B (2019) Scientists rise up against statistical significance. *Nature* 567(7748):305-307.

23. Claridge-Chang A & Assam PN (2016) Estimation statistics should replace significance testing. *Nat Methods* 13(2):108-109.
24. Ho J, Tumkaya T, Aryal S, Choi H, & Claridge-Chang A (2019) Moving beyond P values: data analysis with estimation graphics. *Nat Methods*.
25. Motulsky HJ & Brown RE (2006) Detecting outliers when fitting data with nonlinear regression - a new method based on robust nonlinear regression and the false discovery rate. *BMC Bioinformatics* 7:123.
