## Supplemental Table 1 for "NAD^+^ repletion rescues female fertility during reproductive ageing"

*Bertoldo* et al 2019

**Supplementary Table 1: 95% confidence intervals for data presented in main figures**

| | |
|---|---|
| Fig. 1a | The unpaired mean difference between 4 and 6 is -0.869 [95.0%CI -1.34, -0.508].<br>The two-sided *P* value of the Mann-Whitney test is 0.0369.<br><br>The unpaired mean difference between 4 and 12 is -1.13 [95.0%CI -1.55, -0.853].<br>The two-sided *P* value of the Mann-Whitney test is 0.0369.<br><br>The unpaired mean difference between 4 and 18 is -1.24 [95.0%CI -1.66, -0.97].<br>The two-sided *P* value of the Mann-Whitney test is 0.0369.<br><br>The unpaired mean difference between 4 and 24 is -1.39 [95.0%CI -1.79, -1.15].<br>The two-sided *P* value of the Mann-Whitney test is 0.0369. |
| Fig. 1b | The unpaired mean difference between Control and NMN is 93.0 [95.0%CI 29.2, 1.7e+02].<br>The two-sided *P* value of the Mann-Whitney test is 0.272. |
| Fig. 1d | The unpaired mean difference between Aged and Aged<br>NMN is 0.0177 [95.0%CI 0.00526, 0.0333].<br>The two-sided *P* value of the Mann-Whitney test is 0.119.<br><br>The unpaired mean difference between Aged and Young is 0.0216 [95.0%CI 0.00935, 0.0347].<br>The two-sided *P* value of the Mann-Whitney test is 0.0119 |
| Fig. 2a | $\chi^2$ test p=0.0503 |
| Fig. 2b | The unpaired mean difference between ctrl and NMN is 3.04 [95.0%CI 0.513, 5.67].<br>The two-sided *P* value of the Mann-Whitney test is 0.0221. |
| Fig. 2c | The unpaired mean difference between Aged and Aged NMN is 1.91 [95.0%CI 1.13, 3.07].<br>The two-sided *P* value of the Mann-Whitney test is 0.00227.<br><br>The unpaired mean difference between Aged and Young is 13.9 [95.0%CI 9.03, 18.8].<br>The two-sided *P* value of the Mann-Whitney test is 0.00477. |
| Fig. 2d | The unpaired mean difference between HFD and HFD<br>NMN is 5.9 [95.0%CI 2.5, 9.4].<br>The two-sided *P* value of the Mann-Whitney test is 0.0122.<br><br>The unpaired mean difference between HFD and chow is 3.1 [95.0%CI 0.1, 5.6].<br>The two-sided *P* value of the Mann-Whitney test is 0.0866. |
| Fig. 2e | The unpaired mean difference between WT and Nmnat1Tg/+ is 2.23 [95.0%CI 0.45, 4.53].<br>The two-sided *P* value of the Mann-Whitney test is 0.0785. |
| Fig. 2f | The unpaired mean difference between WT and Nmnat3Tg/+ is -0.623 [95.0%CI -2.13, 2.34].<br>The two-sided *P* value of the Mann-Whitney test is 0.341. |
| Fig. 2g | The unpaired mean difference between Aged and Aged<br>NMN is 2.67 [95.0%CI 0.509, 4.9].<br>The two-sided *P* value of the Mann-Whitney test is 0.0276.<br><br>The unpaired mean difference between Aged and Young is 4.58 [95.0%CI 2.47, 6.89].<br>The two-sided *P* value of the Mann-Whitney test is 0.000593. |
| Fig. 2h | The paired mean difference between ctrl and NMN is 11.7 [95.0%CI -5.55, 25.9].<br>The two-sided *P* value of the Wilcoxon test is 0.144. |
| Fig. 2i | The unpaired mean difference between 0 and 2 is 2.67 [95.0%CI -0.638, 6.91].<br>The two-sided *P* value of the Mann-Whitney test is 0.575.<br><br>The unpaired mean difference between 0 and 7 is 5.34 [95.0%CI 1.7, 8.72].<br>The two-sided *P* value of the Mann-Whitney test is 0.0185.<br><br>The unpaired mean difference between 0 and 14 is 5.09 [95.0%CI 1.95, 9.55].<br>The two-sided *P* value of the Mann-Whitney test is 0.0209.<br><br>The unpaired mean difference between 0 and 28 is 7.46 [95.0%CI 3.52, 12.7].<br>The two-sided *P* value of the Mann-Whitney test is 0.00855. |

| | |
|---|---|
| Fig. 2j | The unpaired mean difference between ctrl and NMN is 1.75 [95.0%CI 0.417, 3.25].<br>The two-sided *P* value of the Mann-Whitney test is 0.0918. |
| Fig. 3c | The unpaired mean difference between chow ctrl and HFD ctrl is 1.49 [95.0%CI 0.899, 2.04].<br>The two-sided *P* value of the Mann-Whitney test is 0.000636.<br><br>The unpaired mean difference between chow ctrl and HFD NMN is 1.83 [95.0%CI 1.1, 2.71].<br>The two-sided *P* value of the Mann-Whitney test is 0.000731. |
| Fig. 3d | The unpaired mean difference between chow ctrl and chow NMN is 1.1 [95.0%CI 0.247, 1.93].<br>The two-sided *P* value of the Mann-Whitney test is 0.0351.<br><br>The unpaired mean difference between chow ctrl and HFD ctrl is 0.0812 [95.0%CI -0.899, 1.01].<br>The two-sided *P* value of the Mann-Whitney test is 0.805.<br><br>The unpaired mean difference between chow ctrl and HFD NMN is 0.957 [95.0%CI 0.0583, 1.76].<br>The two-sided *P* value of the Mann-Whitney test is 0.0351. |
| Fig. 3e | The unpaired mean difference between chow ctrl and chow NMN is 3.57 [95.0%CI -0.378, 8.07].<br>The two-sided *P* value of the Mann-Whitney test is 0.273.<br><br>The unpaired mean difference between chow ctrl and HFD ctrl is 3.38 [95.0%CI 0.0225, 6.92].<br>The two-sided *P* value of the Mann-Whitney test is 0.109.<br><br>The unpaired mean difference between chow ctrl and HFD NMN is 4.91 [95.0%CI 1.46, 9.65].<br>The two-sided *P* value of the Mann-Whitney test is 0.0351. |
| Fig. 3f | The unpaired mean difference between chow ctrl and chow NMN is -0.6 [95.0%CI -9.6, 7.1].<br>The two-sided *P* value of the Mann-Whitney test is 0.97.<br><br>The unpaired mean difference between chow ctrl and HFD ctrl is -6.1 [95.0%CI -16.8, 3.7].<br>The two-sided *P* value of the Mann-Whitney test is 0.364.<br><br>The unpaired mean difference between chow ctrl and HFD NMN is -15.1 [95.0%CI -26.9, -2.3].<br>The two-sided *P* value of the Mann-Whitney test is 0.0691. |
| Fig. 5a | Day 3: The paired mean difference between ctrl and NMN is 19.0 [95.0%CI -6.9, 39.3].<br>The two-sided *P* value of the Wilcoxon test is 0.225.<br>Day 4: The paired mean difference between ctrl and NMN is 19.5 [95.0%CI 1.33, 34.6].<br>The two-sided *P* value of the Wilcoxon test is 0.0431.<br>Day 5: The paired mean difference between ctrl and NMN is 21.8 [95.0%CI 7.02, 39.4].<br>The two-sided *P* value of the Wilcoxon test is 0.0679.<br>Day 6: The paired mean difference between ctrl and NMN is 31.7 [95.0%CI 14.4, 46.1].<br>The two-sided *P* value of the Wilcoxon test is 0.0277. |
| Fig. 5b | The unpaired mean difference between ctrl and 0.01 is -8.73 [95.0%CI -25.8, 5.97].<br>The two-sided *P* value of the Mann-Whitney test is 0.402.<br><br>The unpaired mean difference between ctrl and 0.1 is 30.9 [95.0%CI 12.0, 47.6].<br>The two-sided *P* value of the Mann-Whitney test is 0.00607.<br><br>The unpaired mean difference between ctrl and 1 is 20.2 [95.0%CI -1.25, 44.2].<br>The two-sided *P* value of the Mann-Whitney test is 0.201.<br><br>The unpaired mean difference between ctrl and 10 is -2.63 [95.0%CI -23.0, 15.0].<br>The two-sided *P* value of the Mann-Whitney test is 0.894. |
| Fig. 5c | The unpaired mean difference between vehicle and 0.001 is -73.7 [95.0%CI -85.7, -64.6].<br>The two-sided *P* value of the Mann-Whitney test is 0.0265.<br><br>The unpaired mean difference between vehicle and 0.01 is -75.9 [95.0%CI -87.9, -68.4].<br>The two-sided *P* value of the Mann-Whitney test is 0.0211. |

| | |
|---|---|
| | The unpaired mean difference between vehicle and 0.1 is -63.2 [95.0%CI -78.9, -43.9].<br>The two-sided *P* value of the Mann-Whitney test is 0.0294.<br><br>The unpaired mean difference between vehicle and 1 is -70.9 [95.0%CI -84.1, -57.9].<br>The two-sided *P* value of the Mann-Whitney test is 0.0265.<br><br>The unpaired mean difference between vehicle and 10 is -75.9 [95.0%CI -87.9, -68.4].<br>The two-sided *P* value of the Mann-Whitney test is 0.0211. |
| Fig. 5d | The unpaired mean difference between FK866 and ctrl is 55.1 [95.0%CI 36.3, 68.6].<br>The two-sided *P* value of the Mann-Whitney test is 1.07e-05.<br><br>The unpaired mean difference between FK866 and FK866 plus NMN (0.1 μM) is 20.7 [95.0%CI 6.64, 32.6].<br>The two-sided *P* value of the Mann-Whitney test is 0.00452.<br><br>The unpaired mean difference between FK866 and FK866 plus NMN (0.5 μM) is 49.6 [95.0%CI 33.8, 63.4].<br>The two-sided *P* value of the Mann-Whitney test is 0.000102.<br><br>The unpaired mean difference between FK866 and FK866 plus NaMN (0.1 μM) is 36.3 [95.0%CI 12.7, 53.0].<br>The two-sided *P* value of the Mann-Whitney test is 0.00199.<br><br>The unpaired mean difference between FK866 and FK866 plus NaMN (0.5 μM) is 45.8 [95.0%CI 23.7, 63.4].<br>The two-sided *P* value of the Mann-Whitney test is 0.000738.<br><br>The unpaired mean difference between FK866 and FK866 plus NaR (0.1 μM) is 53.5 [95.0%CI 30.2, 69.1].<br>The two-sided *P* value of the Mann-Whitney test is 0.000115.<br><br>The unpaired mean difference between FK866 and FK866 plus NaR (0.5 μM) is 71.5 [95.0%CI 53.0, 87.8].<br>The two-sided *P* value of the Mann-Whitney test is 0.00126.<br><br>The unpaired mean difference between FK866 and FK866 plus NR (0.1 μM) is -7.99 [95.0%CI -18.8, 6.73].<br>The two-sided *P* value of the Mann-Whitney test is 0.623.<br><br>The unpaired mean difference between FK866 and FK866 plus NR (0.5 μM) is 58.2 [95.0%CI 27.1, 79.1].<br>The two-sided *P* value of the Mann-Whitney test is 0.00221. |
| Fig. 5e | The unpaired mean difference between ctrl and 0.001 μM is -18.4 [95.0%CI -35.8, -5.22].<br>The two-sided *P* value of the Mann-Whitney test is 0.0416.<br><br>The unpaired mean difference between ctrl and 0.01 μM is -9.14 [95.0%CI -30.3, 8.69].<br>The two-sided *P* value of the Mann-Whitney test is 0.514.<br><br>The unpaired mean difference between ctrl and 0.1 μM is -8.45 [95.0%CI -20.8, 5.06].<br>The two-sided *P* value of the Mann-Whitney test is 0.255.<br><br>The unpaired mean difference between ctrl and 1 μM is -15.0 [95.0%CI -34.3, 2.12].<br>The two-sided *P* value of the Mann-Whitney test is 0.222.<br><br>The unpaired mean difference between ctrl and 10 μM is -13.9 [95.0%CI -26.6, -0.85].<br>The two-sided *P* value of the Mann-Whitney test is 0.0963.<br><br>The unpaired mean difference between ctrl and 50 μM is 10.1 [95.0%CI 2.14, 19.3].<br>The two-sided *P* value of the Mann-Whitney test is 0.186. |

| | |
|---|---|
| | The unpaired mean difference between ctrl and 100 µM is 3.77 [95.0%CI -14.4, 22.0].<br>The two-sided *P* value of the Mann-Whitney test is 0.768.<br><br>The unpaired mean difference between ctrl and FK866<br>(0.1 µM) is -76.0 [95.0%CI -82.5, -66.9].<br>The two-sided *P* value of the Mann-Whitney test is 0.055. |
| Fig. 6a | The unpaired mean difference between ctrl and 1 µM sirt. is -25.9 [95.0%CI -41.0, -10.8].<br>The two-sided *P* value of the Mann-Whitney test is 0.105.<br><br>The unpaired mean difference between ctrl and 10 µM sirt. is -32.7 [95.0%CI -41.2, -18.8].<br>The two-sided *P* value of the Mann-Whitney test is 0.0304.<br><br>The unpaired mean difference between ctrl and 100 µM sirt. is -29.3 [95.0%CI -42.7, -16.1].<br>The two-sided *P* value of the Mann-Whitney test is 0.0304.<br><br>The unpaired mean difference between ctrl and 1 µM split. is -33.9 [95.0%CI -43.0, -24.6].<br>The two-sided *P* value of the Mann-Whitney test is 0.0304.<br><br>The unpaired mean difference between ctrl and 10 µM split. is -32.3 [95.0%CI -47.1, -22.6].<br>The two-sided *P* value of the Mann-Whitney test is 0.0304.<br><br>The unpaired mean difference between ctrl and 100 µM split. is -48.8 [95.0%CI -64.0, -23.0].<br>The two-sided *P* value of the Mann-Whitney test is 0.0304. |
| Fig. 6b | The unpaired mean difference between ctrl and 1 µM sirt. is -14.0 [95.0%CI -17.4, -10.2].<br>The two-sided *P* value of the Mann-Whitney test is 0.000565.<br><br>The unpaired mean difference between ctrl and 10 µM sirt. is -11.7 [95.0%CI -14.6, -8.51].<br>The two-sided *P* value of the Mann-Whitney test is 1.03e-08.<br><br>The unpaired mean difference between ctrl and 100 µM sirt. is -16.8 [95.0%CI -20.2, -13.5].<br>The two-sided *P* value of the Mann-Whitney test is 4.65e-12.<br><br>The unpaired mean difference between ctrl and 1 µM split. is -11.9 [95.0%CI -15.5, -8.65].<br>The two-sided *P* value of the Mann-Whitney test is 3.26e-07.<br><br>The unpaired mean difference between ctrl and 10 µM split. is -12.8 [95.0%CI -15.8, -9.63].<br>The two-sided *P* value of the Mann-Whitney test is 2.05e-09.<br><br>The unpaired mean difference between ctrl and 100 µM split. is -17.6 [95.0%CI -21.5, -13.6].<br>The two-sided *P* value of the Mann-Whitney test is 4.5e-08. |
| Fig. 6c | The unpaired mean difference between ctrl and NMN is 24.0 [95.0%CI 6.13, 45.9].<br>The two-sided *P* value of the Mann-Whitney test is 0.0806.<br><br>The unpaired mean difference between ctrl and sirtinol is -23.9 [95.0%CI -35.5, -11.1].<br>The two-sided *P* value of the Mann-Whitney test is 0.00187.<br><br>The unpaired mean difference between ctrl and sirtinol+NMN is -5.03 [95.0%CI -18.4, 11.7].<br>The two-sided *P* value of the Mann-Whitney test is 0.207. |
| Fig. 6d | The unpaired mean difference between 0 µM and 10 µM is -4.1 [95.0%CI -11.9, 1.23].<br>The two-sided *P* value of the Mann-Whitney test is 0.663.<br><br>The unpaired mean difference between 0 µM and 30 µM is -0.4 [95.0%CI -10.1, 9.27].<br>The two-sided *P* value of the Mann-Whitney test is 1.0.<br><br>The unpaired mean difference between 0 µM and 0 µM + sirtinol is -28.1 [95.0%CI -36.7, -19.5].<br>The two-sided *P* value of the Mann-Whitney test is 0.0809. |

| | |
|---|---|
| | The unpaired mean difference between 0 μM and 10 μM + sirtinol is -4.67 [95.0%CI -16.0, 2.53].<br>The two-sided *P* value of the Mann-Whitney test is 0.663.<br><br>The unpaired mean difference between 0 μM and 30 μM + sirtinol is 1.1 [95.0%CI -7.1, 7.33].<br>The two-sided *P* value of the Mann-Whitney test is 0.663. |
| Fig. 6e | The unpaired mean difference between ctrl and pifithrin 10 is 3.34 [95.0%CI -0.144, 6.74].<br>The two-sided *P* value of the Mann-Whitney test is 0.0521.<br><br>The unpaired mean difference between ctrl and pifithrin 30 is -1.11 [95.0%CI -4.55, 2.16].<br>The two-sided *P* value of the Mann-Whitney test is 0.712.<br><br>The unpaired mean difference between ctrl and sirtinol is -16.8 [95.0%CI -19.8, -14.0].<br>The two-sided *P* value of the Mann-Whitney test is 7.38e-18.<br><br>The unpaired mean difference between ctrl and sirtinol+pifithrin 10 is -2.96 [95.0%CI -6.25, 0.158].<br>The two-sided *P* value of the Mann-Whitney test is 0.134.<br><br>The unpaired mean difference between ctrl and sirtinol+pifithrin 30 is -0.361 [95.0%CI -3.77, 3.03].<br>The two-sided *P* value of the Mann-Whitney test is 0.783. |
